# Crosstalk with keratinocytes causes GNAQ oncogene specificity in melanoma

**DOI:** 10.1101/2021.07.26.453858

**Authors:** Oscar Urtatiz, Amanda Haage, Guy Tanentzapf, Catherine D. Van Raamsdonk

## Abstract

Different melanoma subtypes exhibit specific and non-overlapping sets of oncogene and tumor suppressor mutations, despite a common cell of origin in melanocytes. For example, activation of the Gα_q/11_ signaling pathway is a characteristic initiating event in primary melanomas that arise in the dermis, uveal tract or central nervous system. It is rare in melanomas arising in the epidermis. Here, we present evidence that in the mouse, crosstalk with the epidermal microenvironment actively impairs the survival of melanocytes expressing the GNAQ^Q209L^ oncogene, providing a new model for oncogene specificity in cancer. The presence of epidermal cells inhibited cell division and fragmented dendrites of melanocytes expressing GNAQ^Q209L^ in culture, while they promoted the growth of normal melanocytes. Differential gene expression analysis of FACS sorted epidermal melanocytes showed that cells expressing GNAQ^Q209L^ exhibit an oxidative stress and apoptosis signature previously linked to vitiligo. Furthermore, PLCB4, the direct downstream effector of Gα_q/11_ signaling, is frequently mutated in cutaneous melanoma alongside P53 and NF1. Our results suggest that a deficiency of PLCB4 promotes cutaneous melanomagenesis by reducing GNAQ driven signaling. Hence, our studies reveal the flip side of the GNAQ/PLCB4 signaling pathway, which was hitherto unsuspected. In the future, understanding how epidermal crosstalk restrains the GNAQ^Q209L^ oncogene could suggest novel melanoma therapies.

## INTRODUCTION

*GNAQ* and *GNA11* encode heterotrimeric G protein alpha subunits that are best known for signaling through phospholipase C-beta (PLCB) to release intracellular calcium stores^1^. The somatic substitution of glutamine 209 or arginine 183 in either *GNAQ* or *GNA11* generates constitutive activity of the G protein and is a characteristic early event in uveal melanoma, occurring in up to 90% of cases^2^. These mutations are also frequent in blue nevi and primary melanomas of the central nervous system^2–4^. Activation of the pathway in uveal melanoma drives cell proliferation and stimulates the mitogen activated protein kinase (’MAPK’) pathway through RasGRP3^5^. Gα_q/11_ activation also activates the Hippo pathway through nuclear localization of YAP1 via a Trio-Rho/Rac signaling circuit^6^. On the other hand, Q209 and R183 mutations in either *GNAQ* or *GNA11* are rare in melanocytic neoplasms arising in the epithelium, which are more likely to carry mutations in the MAPK pathway components, *BRAF*, *NRAS* and *NF1*^3, 7, 8^. The underlying mechanism for the restriction of *GNAQ* and *GNA11* mutations to non-epithelial melanomas is unknown.

There are several possible, non-exclusive explanations for this phenomenon. First, an over- representation of mutations in specific genes in certain neoplasms could be related to exposure to different mutagens in different parts of the body. Concerning melanoma, the skin areas exposed to UV radiation via sunlight accumulate many somatic mutations; characteristically, these are C to T transitions^9^. Most UV radiation does not penetrate very deeply into the body and some areas of the body are rarely exposed, which could generate a varying spectrum of mutations^10^.

Secondly, melanocytes arise from neural crest cells all along the anterior-posterior axis. The molecular pathways for melanocyte cell fate determination along this axis are poorly understood^11^. However, there are demonstrated differences in the potential for melanomagenesis of melanocytes arising at different anterior-posterior positions in the mouse embryo^12^. It is possible that the developmental origin of a melanocyte could influence its migration during embryogenesis, its differentiation potential and program long-term gene expression patterns. Therefore, some melanocytes might only respond to specific driver mutations due to intrinsic differences generated during development.

Lastly, different microenvironments surround melanocytes located in different parts of the body, which could impact the effects of mutations. Direct cell-cell contact or paracrine signaling produced by the tissue-specific microenvironment might allow the transformation of cells only with certain mutational events, forcing the selection of specific driver mutations in melanoma^13^. Epidermal melanocytes interact mainly with keratinocytes, while internal melanocytes interact with various mesodermal stromas. Therefore, crosstalk between melanocytes and their cellular neighbors might prevent transformation or proliferation driven by specific signaling pathways.

In this paper, we investigated these possibilities. We forced the expression of oncogenic GNAQ^Q209L^ in all melanocytes in mice beginning in adulthood using *Tyr-creER*. This drove the loss of melanocytes from the interfollicular (IFE) epidermis of the tail, while dramatically increasing melanocyte growth in the dermis. This observation demonstrated that GNAQ^Q209L^ expression was not simply neutral for epidermal melanocytes; it was deleterious. We established primary cultures of normal or GNAQ^Q209L^ expressing melanocytes sorted from the mouse tail epidermis and used time lapse microscopy to quantify melanocyte cell dynamics in the presence of either fibroblasts or keratinocytes. We discovered that paracrine signaling from the epidermis reversibly switches GNAQ^Q209L^ from an oncogene to an inhibitor of melanocyte survival and proliferation. Differential expression analysis of melanocytes sorted from the tail epidermis showed that the melanocytes expressing GNAQ^Q209L^ exhibited alterations in gene expression related to cell adhesion, axon extension, oxidative stress and apoptosis. Since GNAQ^Q209L^ signaling was deleterious to epidermal melanocytes, we wondered whether loss of function mutations in *PLCΒ4* might promote melanomagenesis. We examined 470 cases in the Cancer Genome Atlas (TCGA) skin cutaneous melanoma (SKCM) dataset and found that 21% had a mutation in *PLCB4*. *PLCB4* mutations were enriched in cases with *TP53* and *NF1* mutations. We describe the relationship between mutation load and the combinations of certain gain and loss of function drivers in the dataset.

## RESULTS

### Forced GNAQ^Q209L^ reduces melanocytes in mouse tail inter-follicular epidermis (IFE)

We previously generated a conditional GNAQ^Q209L^ allele wherein oncogenic human GNAQ^Q209L^ is expressed from the *Rosa26* locus in mice following the removal of a loxP-flanked stop cassette (“*R26-fs-GNAQ^Q209L^*”)^14^. We previously showed that forcing GNAQ^Q209L^ expression in melanocytes beginning during embryogenesis using the *Mitf-cre* transgene reduced the number of melanocytes in the inter-follicular epidermis (IFE)^14, 15^. To test whether GNAQ^Q209L^ affected melanocyte establishment or melanocyte maintenance, we used the tamoxifen (TM) inducible melanocyte-specific Cre transgene, *Tyrosinase (Tyr)-creERT^2^* with *R26-fs-GNAQ^Q209L^* to induce GNAQ^Q209L^ expression in melanocytes beginning in adulthood. We also included the *Rosa26- LoxP-Stop-LoxP-LacZ* reporter to label cells expressing GNAQ^Q209L^ with LacZ ^16, 17^. TM was administered once per day for 5 days at 4 weeks of age. One week following the last dose of TM, half of the mice were euthanized to collect and stain tail epidermal sheets with X-gal to determine the average number of melanocytes (LacZ-positive cells) per scale. At this time point, *R26-fs-GNAQ^Q209L^/R26-fs-LacZ*; *Tyr-creERT^2^*/+ (hereafter referred to as “GNAQ-lacZ”) mice and *+/R26-fs-LacZ*; *Tyr-creERT^2^*/+ (hereafter referred to as “WT-LacZ”) mice showed no significant difference in melanocyte numbers (p=0.18, **Figures 1A, B**). However, 8 weeks following the last dose of TM, GNAQ-LacZ mice had on average 50% fewer melanocytes per scale than WT-LacZ mice (p=0.045, **Figures 1A, B)**. While scales in WT-LacZ mice were pigmented evenly, many scales in GNAQ-LacZ mice exhibited partial loss of melanin 8 weeks after TM (p=0.048, **Figure 1C, D**). These results demonstrate that GNAQ^Q209L^ signaling inhibits melanocyte maintenance in the IFE.

**Figure 1.**
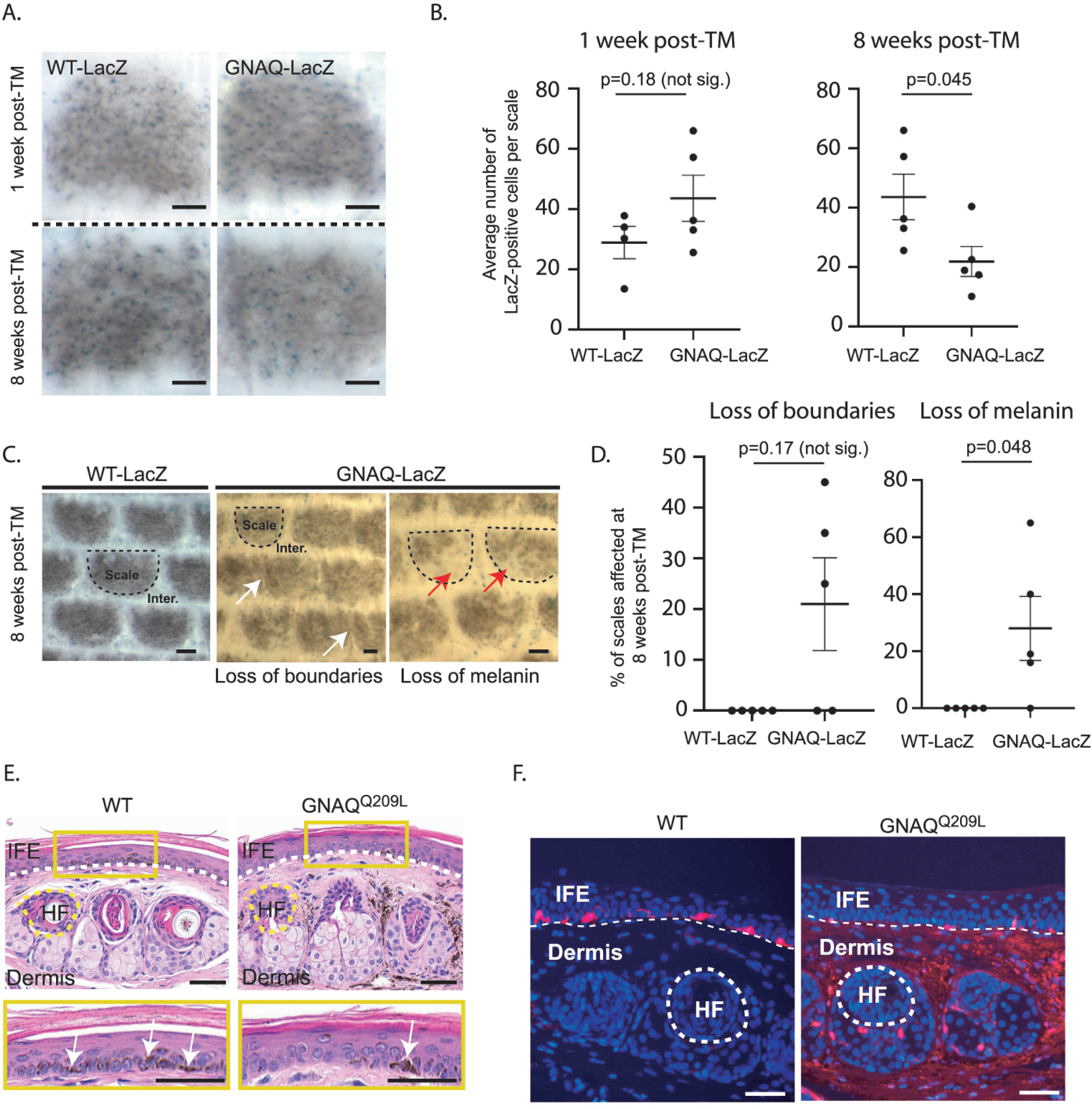
Forced GNAQ-Q209L signaling reduces the number of melanocytes in the IFE. **A)** Representative example of Lael-positive cells within scales of WT-Lacz and GNAQ-LacZ tails at 1 week or 8 weeks post-tamoxifen (TM) treatment. **8)** Quantification of the average number of Lacz-positive cells per scale at 1 week and 8 weeks post-TM treatment. (Each point represents avg for 1 mouse; mean+/- SEM, Unpaired t-test). **C)** Representative examples of X-gal stained whole mount epidermal tail sheets in WT-Lacz and GNAQ-LacZ mice at 8 weeks post-TM, showing loss of boundaries in scale pigmentation (white arrows) or loss of melanin (red arrows) in GNAQ-LacZ mice. Example scales are outlined in dashed line for reference. **0)** Percentage of epidermal scales exhibiting loss of boundaries or loss of melanin in 5 WT-Lacz and 5 GNAQ-LacZ mice at 8 weeks post-TM. (Each point represents% for 1 mouse; Kolmogorov-Smirnov test, mean+/- SEM). **E)** H&E stained cross-sections of tail skin in WT and GNAQ-Q209L mice. The yellow box below shows a magnified area of the inter-follicular epidermis (IFE). Less melanin was observed in the IFE of GNAQ-Q209L skin (white arrows point to examples). Dashed lines indicate the boundaries between the IFE, dermis, and an example hair follicle (HF). **F)** tdTomato expression (red) in cross sections of tail skin of WT and GNAQ-Q209L mice at 4 weeks of age showing a reduced number of melanocytes (tdTomato+ cells) in the IFE of GNAQ-Q209L mice and an abnormal expansion of melanocytes in the dermis. Sections are counterstained with DAPI (blue). Dashed lines indicate the boundaries between the IFE, dermis and an example hair follicle (HF). In A and C, scale bars represent 100 um, while in E and F, scale bars represent 50 um.

### The effects of GNAQ^Q209L^ signaling in melanocytes are reversible

To isolate melanocytes from the epidermis for analysis *in vitro*, we used *Mitf-cre* with *Rosa26- LoxP-Stop-LoxP-tdTomato* and *Rosa26-fs-GNAQ^Q209L^* to label the melanocyte lineage with a robust and intense fluorescent tdTomato signal. To confirm appropriate labeling, we first compared sections of the tail skin of +/*R26-fs-tdTomato; Mitf-cre/+* (“WT”) and *R26-fs- GNAQ^Q209L^/ R26-fs-tdTomato; Mitf-cre/+* (“GNAQ^Q209L^”) mice at 4 weeks of age (**Figure 1E-F**).

Sections of WT tail skin exhibited tdTomato-positive cells in the IFE, located as expected on the basal membrane, as well as some tdTomato-positive cells in the dermis. In contrast, sections of GNAQ^Q209L^ tail skin contained fewer tdTomato-positive cells in the IFE while exhibiting an abnormally extensive tdTomato signal in the dermis.

Next, we isolated tdTomato-positive melanocytes from the IFE of WT and GNAQ^Q209L^ mice by Fluorescent Activated Cell Sorting (FACS) to study the melanocytes in *in vitro* settings. To obtain a single cell suspension for FACS, we first split the tail skin IFE from the underlying dermis. Then, we dissociated the scales within the IFE with trypsin to obtain melanocytes and keratinocytes as dispersed single cells. As expected, fewer tdTomato-positive cells were sorted from GNAQ^Q209L^ mice (0.61%) compared to WT mice (0.99%) (p=0.012, **Figure 2A,B**). Previous studies have estimated that melanocytes account for ∼1.5% of the total cells in the IFE^18^.

**Figure 2.**
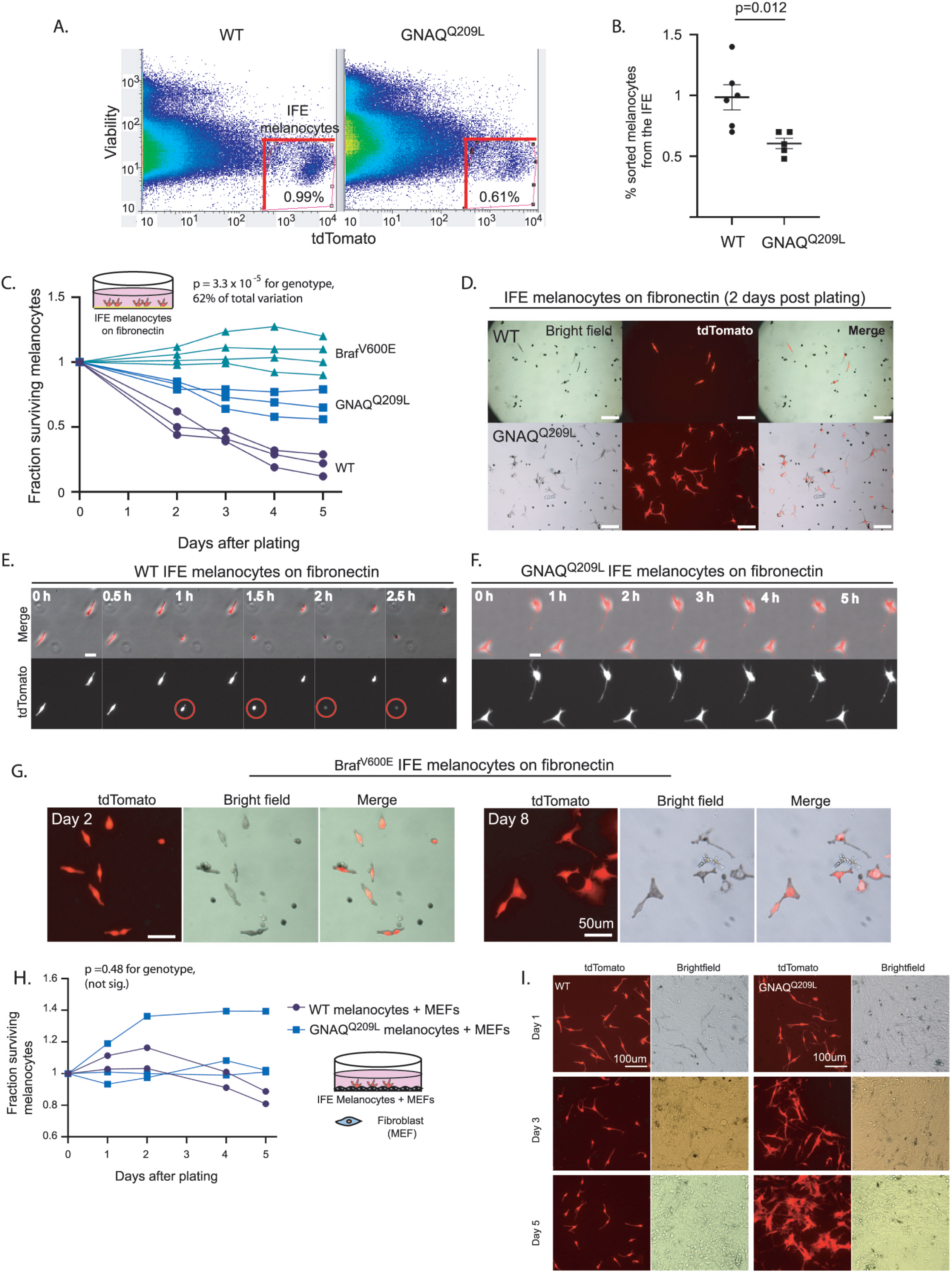
FACS sorted GNAQ-Q209L and BRAFV-600E IFE melanocytes cultured on fibronectin have increased survival compared to WT. **A)** Representative examples of FACS dot plots of single-cell suspension from the IFE epidermis of WT and GNAQ-Q209L mice. **B)** Average percentage of tdTomato+ cells sorted from the IFE of WT and GNAQ-Q209L mice, with a significantly smaller percentage in GNAQ-Q209L. (Each point represents% from 1 mouse, mean ± SEM. Unpaired t test). **C)** Fraction of surviving WT, GNAQ-Q209L, and Braf-V600E FACS sorted melanocytes plated on fibronectin (each line represents an independently derived primary culture, 2 way Anova). **0)** Representative images of melanocyte morphology 2 days post-plating on fibronectin-coated wells, showing increased dendrite formation in GNAQ-Q209L melanocytes. **E)** Time-lapse microscopy of two WT melanocytes plated on fibronectin. The circled cell adopted a round shape shortly before being lost from view. **F)** Time-lapse microscopy of two GNAQ-Q209L melanocytes showing a dendritic cell morphology that remained stable over time. **G)** Representative images of BRAF-V600E melanocytes at 2 and 8 days post-plating on fibronectin. Scale bars represent 100 um in D, 20 um in E and F, and 50 um in G. **H)** Fraction of surviving WT and GNAQ-Q209L IFE melanocytes co-cultured with mouse embryonic fibroblasts (MEFs), showing initial growth above the baseline for both genotypes before loss began (each line represents an independently derived primary culture, 2 way Anova). I) Representative images of FACS sorted WT and GNAQ-Q209L IFE melanocytes co-cultured with MEFs. While the WT and GNAQ-Q209L IFE melanocytes had a similar spindle cell morphology at day 1, the GNAQ-Q209L melanocytes progressively developed large and abnormal shapes.

We then seeded the sorted IFE melanocytes on fibronectin-coated plates to study baseline survival. The GNAQ^Q209L^ expressing melanocytes survived better than the WT melanocytes (p=3.3 x 10^-5^ for genotype, 2 way ANOVA analysis, **Figure 2C**). Hence, the growth-inhibiting effects of GNAQ^Q209L^ expression in IFE melanocytes are reversible. When removed from the epidermis and grown on fibronectin, IFE melanocyte survival was increased by GNAQ^Q209L^ expression.

To compare the effects of GNAQ^Q209L^ with another well studied melanoma oncogene, we used the same methods to obtain Braf^V600E^ expressing melanocytes from the mouse tail IFE. In mice, the expression of Braf^V600E^ leads to melanocytic over-growth in both the tail dermis and epidermis^19^. We crossed the conditional *Braf^V600E^* allele, which expresses *Braf^V600E^* from the endogenous *Braf* locus^20^ to *R26-fs-tdTomato* and *Mitf-cre*. Then, we used FACS to isolate epidermal melanocytes from the tail IFE of 4-week old mice (+/*R26-fs-tdTomato; Mitf-cre/+; Braf^CA^/+,* hereafter referred to as “Braf^V600E^”). We found that Braf^V600E^ expression increased the survival of sorted IFE melanocytes plated on fibronectin, even more so than GNAQ^Q209L^ (**Figure 2C**). Melanocytes expressing Braf^V600E^ could be maintained in culture for at least 8 days (**Figure 2G**).

We observed a striking difference in cell morphology of GNAQ^Q209L^ melanocytes compared to WT melanocytes grown *in vitro*. GNAQ^Q209L^ melanocytes exhibited a more dendritic morphology with several protrusions per cell, while WT melanocytes appeared mostly spindle- shaped (**Figure 2D**). To study the cellular dynamics of the melanocytes, we analyzed time lapse microscopy for 20 hours (**Figure 2E, F**). No migration, formation of new protrusions or cell divisions were observed in the melanocytes of either the WT or *GNAQ^Q209L^* cultures plated on fibronectin alone. However, in WT cultures, some melanocytes were observed to adopt a round shape, after which the cells were immediately lost from view (presumed cell death, example shown in the circled cell in **Figure 2E**).

As another test of GNAQ^Q209L^ reversibility, we isolated mouse embryonic fibroblasts (MEFs) and plated them onto fibronectin-coated plates to simulate a dermis-like microenvironment. Then we added FACS sorted IFE melanocytes from either WT or GNAQ^Q209L^ mouse tails. The growth of both WT and GNAQ^Q209L^ expressing melanocytes was stimulated by MEF co-culture, as there was an increase in cell number above the original number plated, unlike on fibronectin alone (**Figure 2H**). In this situation, we did not detect a difference in survival between WT and GNAQ^Q209L^ melanocytes (p=0.48, 2 way ANOVA). When grown with MEFs, GNAQ^Q209L^ melanocytes grew much larger and irregularly shaped than WT melanocytes (**Figure 2I**), similar to their appearance in the dermis of tail sections (**Figure 1F**).

To summarize, our findings suggest that the attrition of GNAQ^Q209L^ expressing IFE melanocytes from the mouse tail epidermis is not due to an inherent characteristic in IFE melanocytes because the phenotype is reversible by switching the microenvironment.

### The IFE impairs the survival and proliferation of GNAQ^Q209L^ melanocytes *in vitro*

Keratinocytes are known to affect melanocyte proliferation and survival in the IFE in a normal context, so we suspected that interactions between keratinocytes and melanocytes in the IFE could be modifying the outcome of GNAQ^Q209L^ signaling. To test this, we dissociated tail IFE into single cells and plated the suspension onto fibronectin-coated plates. Co-culturing with dispersed IFE provided a significant boost to WT melanocyte survival compared to fibronectin coating alone (p=0.0071 for culture condition, 2 way ANOVA, **Figure 3A**). In contrast, the co-culture of IFE with GNAQ^Q209L^ expressing melanocytes caused a significant reduction in melanocyte survival throughout the five days of the experiment (p=0.0053 for culture condition, 2 way ANOVA, **Figure 3B**).

**Figure 3.**
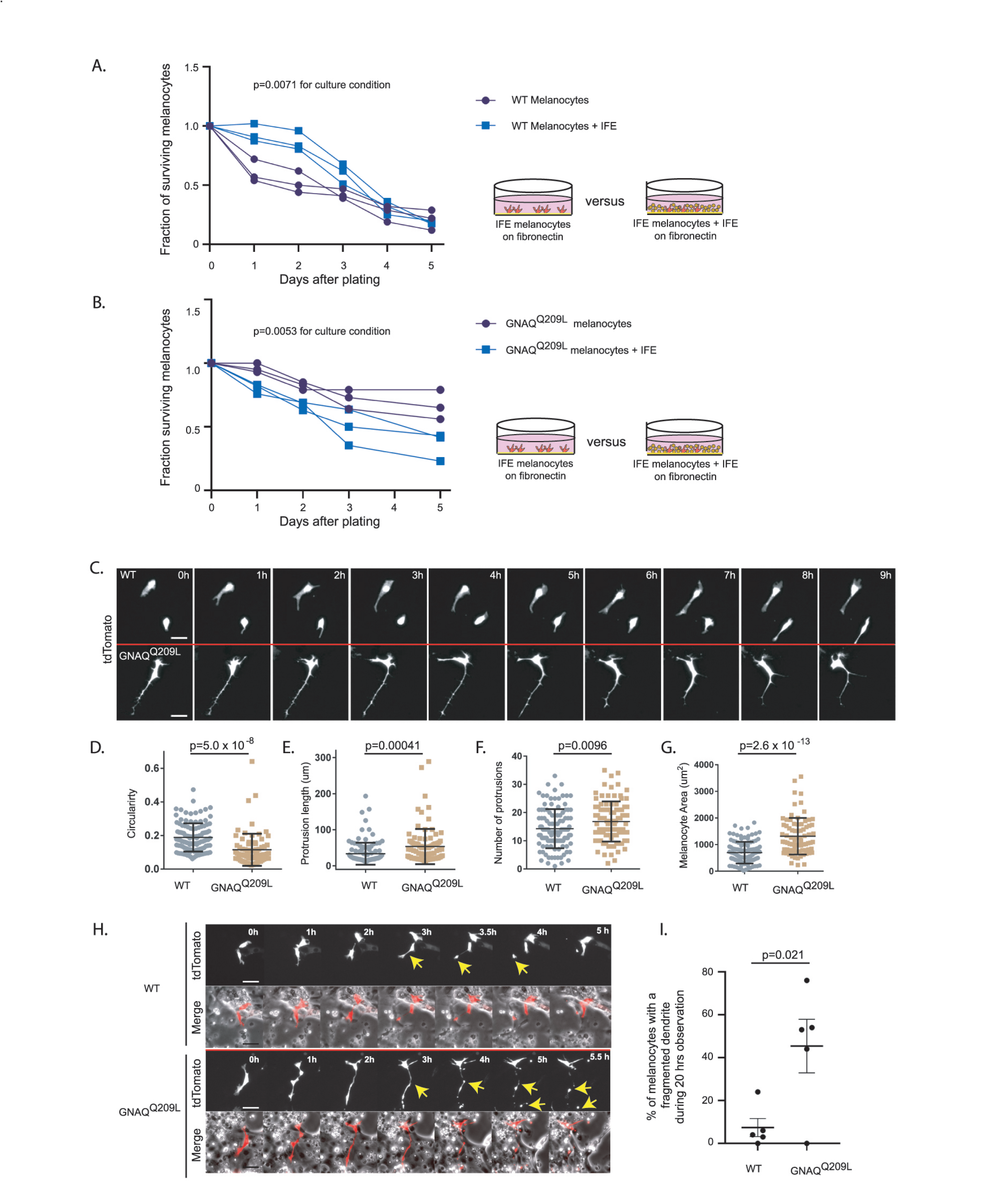
The IFE impairs survival and alters pseudopod dynamics in GNAQ-Q209L melanocytes. **A)** Survival of unsorted WT melanocytes plated with its IFE, compared to sorted IFE WT melanocytes plated onto fibron ectin. The presence of IFE increased the survival of WT melanocytes. (Each line represents one indepedent ly derived culture, 2 way Anova). **B)** Survival of unsorted GNAQ-Q209L melanocytes plated with its IFE, compared to sorted IFE GNAQ-Q209L melanocytes plated onto fibronectin. The presence of IFE decreased t he survival of GNAQ-Q209L melanocytes. (Each line represents one indepedently derived culture, 2 way Anova). **C)** Time-lapse images show representative WT and GNAQ-Q209L melanocytes co-cultured with IFE between 0 and 8 hours. The GNAQ-Q209L cell exhibits abnormally long dendrites and a less circular (more polygonal) cell body shape. **0-G)** Quantification of ci rcularity (D), protrusion length (E), number of protrusions (F), and melanocyte area (G), in WT and GNAQ-Q209L melanocytes co-cultured with IFE. (Each point represents the mesurement of one cell, mean ± SEM; Unpaired t test). **H)** Time-lapse microscopy showing a representative example of dendrite fragmentation in WT and GNAQ-Q209L melanocytes.Arrows indicate dendrite breakage points and subsequent fragments that form up into balls. I) Quant ification of the percent of cells experiencing dendrite fragmentation in WT and GNAQ-Q209L melanocytes cultured with IFE. (Each point represents the mesurement from one culture, mean ± SEM; Unpaired t test). Scale bars represent 40 um in C and H.

Observation using time lapse microscopy showed that both WT and GNAQ^Q209L^ melanocytes exhibited different behavior when cultured with IFE compared to culture on fibronectin alone, with both WT and GNAQ^Q209L^ melanocytes migrating and extending and retracting protrusions (**Figure 3C**, example **Videos 1 and 2**). GNAQ^Q209L^ expressing melanocytes exhibited longer and more numerous protrusions (p=0.0096 and p=0.00041, respectively, **Figure 3E, 3F**). Perhaps related to the unusually long protrusion length, breakage and fragmentation of dendrites was observed in 45% of the GNAQ^Q209L^ melanocytes tracked by time lapse microscopy for 20 hours (**Figure 3H, 3I**). In comparison, this phenomenon was observed in just 5% of WT melanocytes (p=0.021). The average cell area of GNAQ^Q209L^ melanocytes was 2.7-fold greater than WT melanocytes (p=2.6x10^-13^, **Figure 3G**) and the cells were less circular (p=5.0x10^-8^, **Figure 3D**). These changes in cell morphology and dynamics did not seem to affect cell migration (**Figure 4A**). We found no difference in the total length traveled or the straightness of path in GNAQ^Q209L^ versus WT melanocytes cultured with IFE (p=0.86 and p=0.18, **Figure 4B, 4C**).

**Figure 4.**
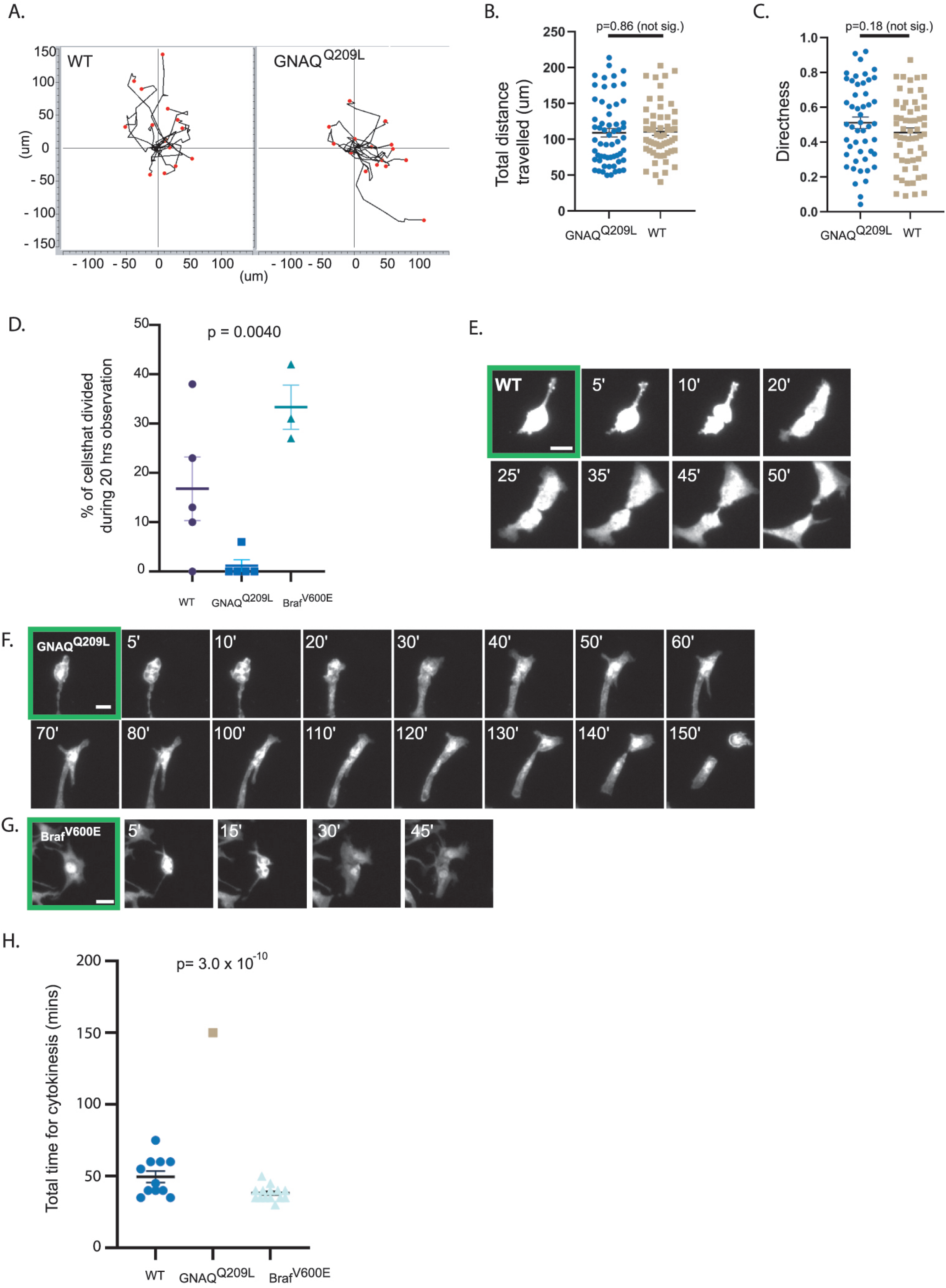
The IFE impairs cell division of GNAQ-Q209L melanocytes. **A)** Migration plots of WT and GNAQ-Q209L melanocytes co-cultured with IFE over 20 hours. **B)** Quantification of total distance traveled in 20 hours. (Each point represents the measurement from 1 cell, mean ± SEM; Unpaired t test). **C)** Quantification of the directness of cell trajectories over 20 hours. Directness=1 is a straight line cell trajectory. (Each point represents the measurement from 1 cell, mean ± SEM; Unpaired t test). **D)** Percentage of tracked cells undergoing division during 20 hours of time-lapse microscopy, when co-cultured with IFE. (Each point represents the measurement from 1 culture, mean ± SEM; Ordinary 1 way AN OVA). **E-G)** Representative examples of cell division events from cleavage furrow formation to separation of daughter cells in a WT melanocyte (in E), a GNAQ-Q209L melanocyte (in F), and a BRAF-V600E melanocyte (in G). **H)** Quantification of the time taken between cleavage furrow formation to separation into daughter cells in melanocytes co-cultured with IFE. (Each point represents the measurement from 1 cell, mean± SEM, Ordinary 1 way ANOVA.) Scale bar in E-G represents 20 um.

We also monitored for cell division events using time lapse microscopy. During the 20 hours of observation, 16% of WT melanocytes co-cultured with IFE underwent a cell division event (**Figure 4D**, example shown in **Figure 4E, video 1**). WT melanocytes smoothly progressed from cleavage furrow formation to the separation of daughter cells and on average this took about 50 minutes (**Figure 4H**). In contrast, only one of the 99 tracked GNAQ^Q209L^ melanocytes divided (p=0.0040, **Figure 4D**). In this cell, furrow formation was not as clear and the process took 150 minutes (**Figure 4F**). We used the same methods to examine the effects of the Braf^V600E^ oncogene on proliferation (**Figure 4G**). During 20 hours of observation, 32% of Braf^V600E^ melanocytes underwent a cell division event (**Figure 4D**). Cell division took the least time in Braf^V600E^ expressing melanocytes (**Figure 4H**).

To determine whether the reduced survival of GNAQ^Q209L^ expressing melanocytes co-cultured with IFE was due to direct contact inhibition or a paracrine mechanism, we used a transwell assay in which we seeded FACS collected IFE melanocytes on a permeable membrane with dissociated IFE from the same animal plated in the well underneath. If direct cell-cell contact was necessary for the inhibitory effect of the IFE, then we would expect increased survival of GNAQ^Q209L^ melanocytes in this culture system compared to simple co-culture. We found that it made no difference whether the IFE was directly in contact with the GNAQ^Q209L^ expressing melanocytes or present in the same dish, separated by a membrane (**Figure 5B**). In WT melanocytes, there was no significant difference in survival for the first three days, after which the transwell melanocytes developed an advantage (**Figure 5A**). We conclude that paracrine signaling from the microenvironment helps switch GNAQ^Q209L^ from an oncogene to an inhibitor of cell survival.

**Figure 5.**
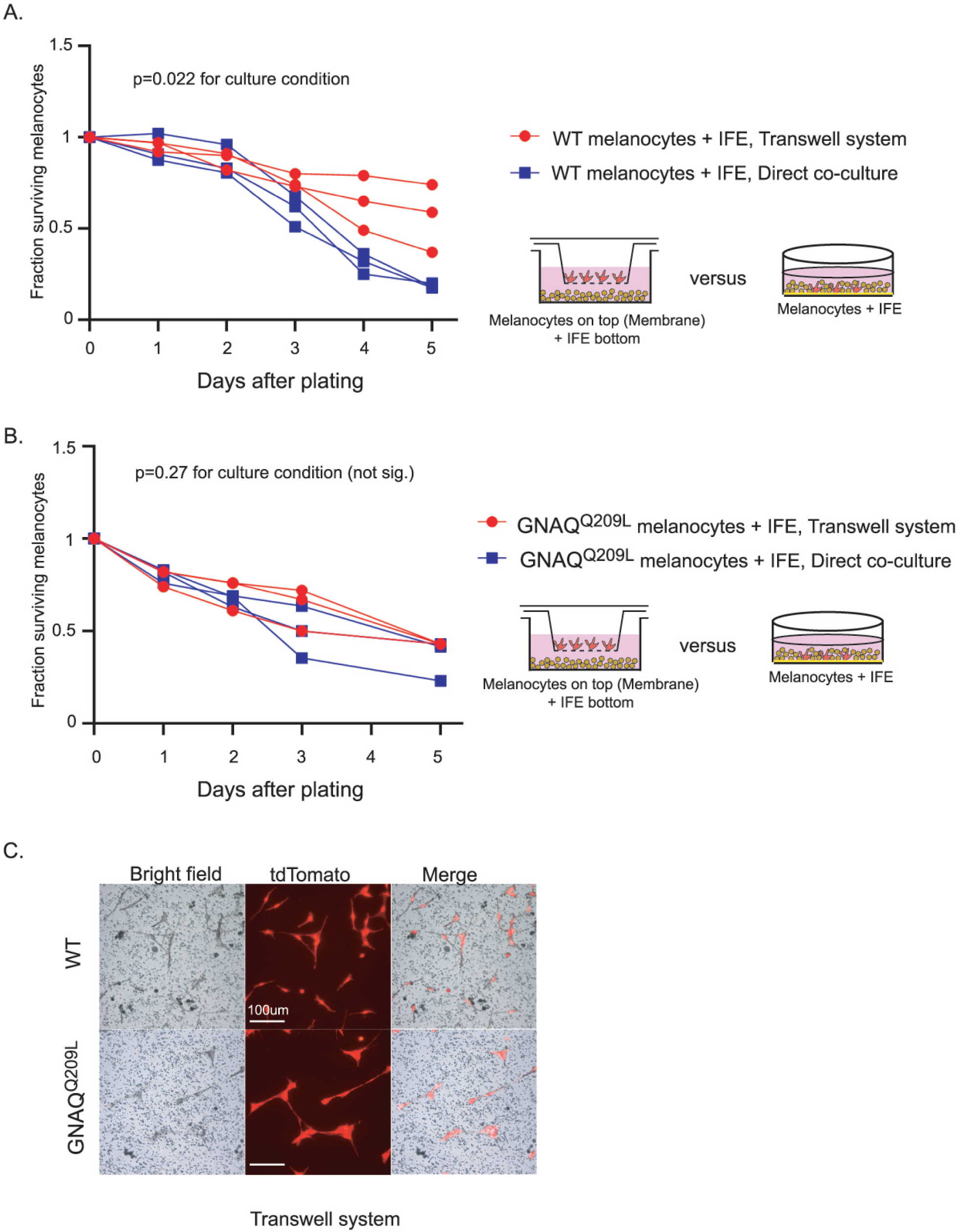
lFE control of melanocytes was maintained in a transwell culture system. **A)** Comparison of survival for WT melanocytes in direct contact with IFE (“direct co-culture”) versus a transwell system where the two populations are separated by a permeable membrane. In WT melanocytes, there was no significant difference in survival for the first three days, after which the transwell melanocytes developed an advantage (Each line represents 1 independent culture, mean ± SEM; 2 way ANOVA.) **B)** Comparison of survival for GNAQ-Q209L melanocytes as in A. There was no significant difference in survival throughout the culture period. (Each line represents 1 independent culture, mean ± SEM; 2 way A NOVA.) **C)** Representative images of WT and GNAQ-Q209L melanocytes at day three on the transwell membrane (no direct cell contact with IFE). GNAQ-Q209L melanocytes were larger than WT melanocytes, as in direct co-culture. Scale bar represents 100 um in C.

Altogether, the cell culture experiments revealed that IFE co-culture stimulates survival and cell division of WT melanocytes, but inhibits these processes in GNAQ^Q209L^ melanocytes. Furthermore, the IFE stimulated protrusion activity in both WT and GNAQ^Q209L^ cells, with the GNAQ^Q209L^ expressing cell hyper-responding. Since both WT and GNAQ^Q209L^ melanocytes were static when plated on fibronectin alone, these experiments show that the IFE plays a vital role in melanocyte regulation. We conclude that crosstalk between melanocytes and the IFE reversibly switches GNAQ signaling from promoting to inhibiting melanocyte growth and survival.

### Gene expression analysis: Regulation of cell adhesion and pseudopod dynamics

To identify the pathways that change in response to GNAQ^Q209L^ expression in IFE melanocytes, we performed RNA sequencing (RNA-seq) immediately after sorting WT and GNAQ^Q209L^ melanocytes from tail IFE (n=6 mice and n=3 libraries per genotype). There were 14,461 genes with an average FPKM > 0.1 in WT and/or GNAQ^Q209L^ melanocytes (**Supplementary Table 1**). Seven genes previously shown to be directly involved in pigment production and melanosome biology were on the list of the top 20 most highly expressed genes in WT melanocytes, validating our melanocyte isolation protocol (**Supplementary Table 2**). These genes were *Pmel*, *Dct*, *Tyrp1*, *Mlana*, *Cd63*, *Slc24a5,* and *Gpnmb*. We analyzed differential gene expression using DEseq2. 1,745 genes were found to be differentially expressed (DE), of which 729 genes were down-regulated and 1016 genes were up-regulated in GNAQ^Q209L^ melanocytes (q-value < 0.05) (**Supplementary Tables 3 and 4**).

We first used the DAVID Functional Annotation Tool (6.8) to understand the biological significance of the DE genes in GNAQ^Q209L^ melanocytes. We analysed the combined list of up and down-regulated DE genes with a log_2_ fold change (“LFC”) of > 2.0 or < -2.0. Using Gene Ontology (GO) analysis and KEGG pathway analysis, there were two dominant signatures. One overlapping set of genes supported the terms: cell adhesion, focal adhesion, extracellular matrix-receptor interactions and extracellular matrix structural constituents (**Figure 6A**, **Supplementary Table 5**). Overlapping genes supported a second signature for the terms: axons, axon guidance and nervous system development (**Supplementary Table 6**).

**Figure 6.**
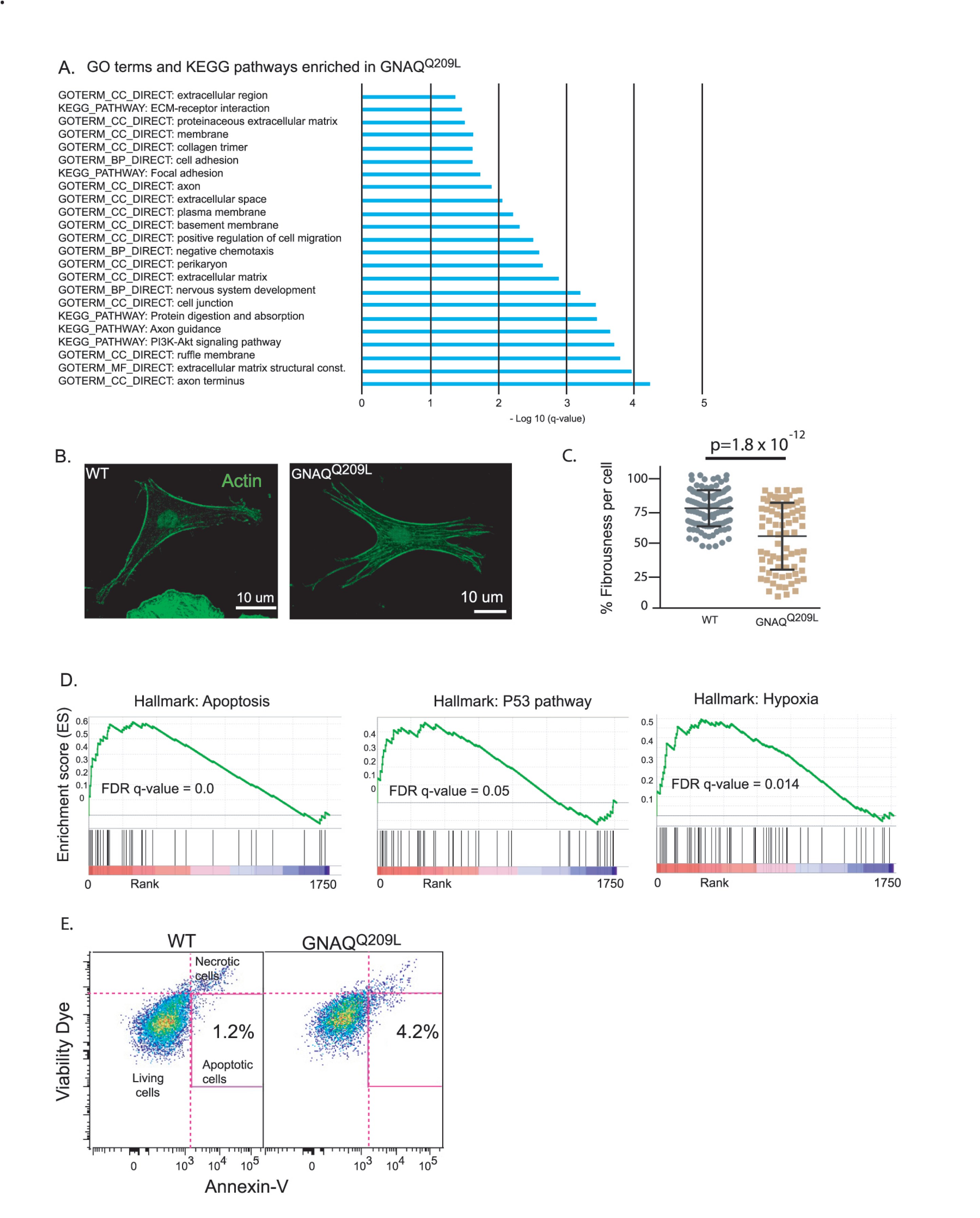
Analysis of GNAQ-Q209L expressing melanocytes in the IFE reveals alterations in the actin cytoskeleton and cell death. **A)** Significant terms identified by gene ontology analysis for DE genes (LFC >2 or <-2) in GNAQ-Q209L melanocytes. **B)** Representative examples of phalloidin staining for f-actin in WT and GNAQ-Q209L melanocytes co-cultured with IFE. **C)** Quantification of fibrousness as a measure of organization of the actin cytoskeleton. (Each point represents a measurement from 1 cell, mean± SEM; Unpaired t test). **D)** Gene set enrichment plots for apoptosis, P53 pathway, and hypoxia hallmarks enriched in GNAQ-Q209L melanocytes. **E)** Percentage of (tdTomato+/Annexin-V collected by FACS from 4 week old tail IFE. N=2 WT and 2 GNAQ-Q209L mice, melanocytes pooled.

For changes related to cellular adhesion, almost all the identified genes were up-regulated in expression. This included types *IV, V*, and *XXVII collagens, Fibronectin 1 (Fn1)*, *Integrin alpha 1, Integrin beta 3, Amigo1, Mcam*, *Cadherin 6, Cercam* and *Esam*. Examining the complete list of DE genes (with no cut-off), we also found that two Rho GEFs that interact with Gα_q/11_, *Trio* and *Kalirin (Kalrn)*, were up-regulated (LFC 0.5 and 1.8). RhoGEFs catalyze GDP to GTP exchange on small Rho guanine nucleotide-binding proteins, regulating the actin cytoskeleton. We also noticed that several genes that play an essential role in lamellipodia formation^21^ including *RhoB*, *RhoC*, *Cdc42*, and *Rock* were all up-regulated by LFC > 1.0. Increased adhesion and cell-ECM interactions could underlie the changes in cell shape and increased cellular area in GNAQ^Q209L^ melanocytes co-cultured with IFE (**Figure 3**).

To examine the actin cytoskeleton in GNAQ^Q209L^ and WT melanocytes, we measured the overall level of actin alignment, termed cell fibrousness, using an *in vitro* quantitative approach previously described by Haage *et al*^22^. First, WT and GNAQ^Q209L^ tdTomato-positive melanocytes co-cultured with IFE were stained for f-actin using phalloidin (**Figure 6B**). Then, confocal z-projections of individuals melanocytes were obtained and processed using quantitative image analysis to determine cell fibrousness. GNAQ^Q209L^ expressing melanocytes exhibited less cell fibrousness than WT melanocytes (p=1.8x10^-12^, **Figure 6C**). Hence, GNAQ^Q209L^ expressing melanocytes have less organized actin cytoskeletons.

Melanocytes are known to share signaling pathways with neurons^23^. The DE genes related to axons (identified by pathway analysis in **Supplementary Table 6**) are diverse in function; however, the number of *Semaphorin* genes on the list stands out, suggesting that these signaling molecules might be important. While the autocrine effects of Semaphorins on melanocytes has not been addressed before, it is known that keratinocytes express semaphorins to regulate melanocytes^24^. Semaphorin signaling controls the formation of cellular extensions such as axons and dendrites in both neurons and melanocytes^25^. Semaphorins bind to two main classes of receptors, plexins and neuropilins. Some semaphorins can also bind the Integrin beta 3 receptor^24^. Scott *et al.* showed that Sema7a binding to Integrin beta 3 induced melanocyte spreading and dendricity, while Sema7a binding to plexin C1 inhibited dendrite formation^24^. Another study found that the absence of Sema3A leads to defasciculation and abnormal extension of certain cranial nerves^26^.

Among the complete list of DE genes, the semaphorins that were up-regulated were *Sema3c, Sema3d, Sema3g, Sema4c* and *Sema4f*, while *Sema3a, Sema3b, Sema4b, Sema5a, Sema6d* and *Sema7a* were down-regulated. *Sema4f* was the most highly up-regulated *Sema* family member (LFC 4.4), followed by *Sema3g* (LFC 2.8). In GNAQ^Q209L^ melanocytes, the expression of *Integrin beta 3* was also up-regulated (LFC 3.6), while *Plexin C1* was down-regulated (LFC - 0.7). These expression changes in *Sema3A*, *Sema7A*, *Integrin beta 3* and *Plexin C1* are consistent with the abnormally long dendrites and increased cellular area in GNAQ^Q209L^ expressing melanocytes^24^.

To summarize, the RNAseq data suggests that the impact of GNAQ^Q209L^ expression on IFE melanocyte morphology is mediated through increased cell adhesion, a disorganized actin cytoskeleton and the promotion of dendrite extensions possibly stimulated by changes in semaphorin signaling.

### Gene expression analysis: Cellular stress and apoptosis

Using the RNAseq data, Gene set enrichment analysis (GSEA) revealed that GNAQ^Q209L^ IFE melanocytes express a pattern of genes related to cellular stress and apoptosis. GSEA showed that GNAQ^Q209L^ melanocytes are enriched in the hallmarks for “apoptosis”, “p53 pathway”, and “hypoxia” (**Figure 6D**). Among the top-ranked DE genes was *Stanniocalcin* (*Stc1*) which encodes a glycoprotein hormone involved in calcium/phosphate homeostasis. *Stc1* was up- regulated by a LFC of 7.1 in GNAQ^Q209L^ melanocytes. Significant up-regulation of Stc1 has been reported in tumors under hypoxic or oxidative stress^27–31^. Furthermore, Stc1 expression down- regulates pro-survival ERK1/2 signaling and reduces the survival of MEFs under conditions of oxidative stress^28^. *Cdkn2a* and *Ccnd1* were up-regulated in GNAQ^Q209L^ expressing melanocytes by a LFC of 5 and 1.7, respectively, similar to the intracellular response to oxidative stress previously described in melanocytes in the disease, vitiligo, which results in the loss of melanocytes from the epidermis^32–34^. Moreover, the *S100A8, S100A9* and *BNIP3* genes, involved in cell death and autophagy via ROS-mediated crosstalk between mitochondria and lysosomes, were all up-regulated in GNAQ^Q209L^ melanocytes by a LFC of 0.7 to 1.7^35^.

In terms of apoptosis, *S100b*, a negative regulator of p53 in melanocytes, was down-regulated by (LFC -2.6), while *Annexin V* (*Anxa5*), a marker of early apoptosis, was up-regulated (LFC 0.6). We also found up-regulation of the pro-apoptotic *Bok* (LFC 1.3), an effector of mitochondrial outer membrane permeabilization (MOMP) that triggers membrane permeabilization and apoptosis in the absence of Bax and Bak^36^. *Cystatin C* (*Cst3*), a major inhibitor of cathepsins^37^, was up-regulated (LFC 2.3) in GNAQ^Q209L^ melanocytes. In response to DNA damage, activated p53 can induce *Cystatin C* expression by binding to regulatory sequences in the first *Cst3* intron. Hence, we hypothesized that cellular stress was stimulating increased apoptosis of GNAQ^Q209L^- expressing melanocytes. To assess the amount of cell death *in vivo*, we dissociated IFE from mouse tails and stained the cells for Annexin-V, which labels cells in early apoptosis. Using FACS, we quantified the percentage of tdTomato and Annexin-V double-positive cells (**Figure 6E**). There was a greater percentage of Annexin-V positive melanocytes from GNAQ^Q209L^ tail epidermis (4.2% versus 1.2%). This supported the GSEA data that suggests that the loss of GNAQ^Q209L^ expressing melanocytes from the IFE involves cellular stress and apoptosis.

### Low frequency of *GNAQ* and *GNA11* oncogenic mutations in human cutaneous melanoma

Our results suggested that oncogenic mutations in *GNAQ* and its closely related homolog, *GNA11,* are rarely found in cutaneous melanoma because the epidermal microenvironment causes Gα_q/11_ signaling to inhibit melanocytes. To determine the frequency of these mutations, we searched the COSMIC database to identify all cutaneous melanoma cases with a *GNAQ* or *GNA11* oncogenic mutation at either Q209 or R183. 2753 and 2295 samples meeting our inclusion criteria (see legend of **Supplementary Tables 7, 8)** had *GNAQ* or *GNA11* mutation status reported, respectively. A total of 23 cases carried an oncogenic mutation in either *GNAQ* (n=11) or *GNA11* (n=12). Five of these cases were actually specified as malignant blue nevus or uveal melanoma in their original publications and so were not cutaneous melanomas. Eight other cases carried one or more mutations in genes also found in uveal^38, 39^ or mucosal^40, 41^ melanoma, but not in cutaneous melanoma: *BAP1*, *SF3B1* or *EIF1AX* and hence were similarly suspect as misclassified. One case was from a cell line and in another case, the frequency of an odd *GNAQ^Q209H^* mutation in the tumor was less than 5%. This left eight cases that we think could have arisen in the epidermis, for an incidence of 0.15% (4/2753) for *GNAQ* and 0.17% (4/2295) for *GNA11*. Details on two of these cases can be found in **Supplementary Table 9**.

### Mutations in *PLCB4* in human cutaneous melanoma

*Phospholipase C beta 4* (*PLCB4)* encodes a protein that binds to Gα_q_ and Gα_11_ and serves as the primary effector of signaling to downstream components for q class heterotrimeric G proteins.

*PLCB4* was linked to Gα_q/11_ signaling in melanoma when a recurrent gain-of-function substitution (*PLCB4^D630Y^*) was found to be mutually exclusive with *GNAQ* and *GNA11* mutations in uveal and CNS melanomas^42, 43^. If Gα_q/11_ signaling inhibits melanocyte growth in the epidermis, we wondered whether loss of function mutations in *PLCΒ4* might promote melanomagenesis. In 2012, Wei *et al.* reported that *PLCB4* was one of the most frequently mutated genes in a study of 52 cutaneous melanomas (freq = 15%)^44^. However, in the TCGA-SKCM study published in 2015, *PLCB4* was not identified as a Significantly Mutated Gene (SMG) by the MutSig and InVEx algorithms used to analyze the sequencing data^45^. These particular bioinformatics methods, while effective, left out some known melanoma driver genes identified by other studies^46^. Hayward *et al.* subsequently reported that *PLCB4* was targeted by loss-of-function gene fusion events in their cutaneous melanoma set^40^.

We re-examined the data concerning *PLCB4* in the TCGA-SKCM dataset, which currently consists of 470 cases (a list of these cases with description can be found in **Supplementary Table 9**). Most cases (n=420) had a primary diagnosis of ‘Malignant melanoma, not otherwise specified (NOS)’. 50 cases were variously described as ‘Epithelioid cell melanoma’, ‘Nodular melanoma’, ‘Amelanotic melanoma’, ‘Lentigo maligna melanoma’, ‘Acral lentiginous melanoma, malignant’, ‘Superficial spreading melanoma’, ‘Spindle cell melanoma, NOS’, or ‘Desmoplastic melanoma, malignant’. Excluding synonymous alterations, small somatic mutations in the exons, introns, UTRs or splice sites of *PLCB4* were found in 97 TCGA-SKCM cases (21% overall). Of the affected cases, 72 carried missense mutations. Other types of mutations (stop gained, splice site, intronic, 5’UTR, 3’UTR and frameshift) occurred along with a missense mutation in 17 additional cases and without one in 8. The most frequent recurrent mutations were S670L, G876E and E1104K, which together were found in 12% of the 97 *PLCB4* affected cases. No case carried a mutation affecting the D630 residue previously identified as a gain-of-function alteration in uveal and CNS melanomas^42, 43^.

We compared *PLCB4* to three of the most frequently mutated tumor suppressor SMGs in melanoma: *PTEN*, *TP53* and *NF1*, considering small somatic mutations within each gene, excluding synonymous alterations (all mutations are listed in **Supplementary Table 9**). The *NRAS^Q^*^61^ and *BRAF^V6^*^00^ oncogenic mutations occur in an exact mutually exclusive pattern in the TCGA-SKCM dataset (116 and 205 cases, respectively). Therefore, we began by comparing the frequency of *PLCB4, PTEN*, *TP53* and *NF1* mutations in these subsets. *PLCB4* was mutant in 24% of the *NRAS^Q^*^61^ mutant cases, 17% of the *BRAF^V6^*^00^ mutant cases and 23% of the remaining cases (hereafter referred to as ‘other’). *TP53* and *NF1* were also less frequent in *BRAF^V6^*^00^ mutant cases and more frequent in the *NRAS^Q^*^61^ and other cases (**Figure 7A**). *NF1* mutations were highly enriched in the ‘other’ subset, as expected from the literature^8, 40^. *PTEN* mutations, in contrast, were more likely to be found in the *BRAF^V6^*^00^ mutant subset compared to the *NRAS^Q^*^61^ mutant subset or other cases. We calculated the average total number of all gene associated small somatic mutations in each of the three subsets and found that the ‘other’ subset had 1.7-fold greater mutation load than the *NRAS^Q^*^61^ mutant subset. Hence, *TP53* and *PLCB4* mutations in the *NRAS^Q^*^61^ subset could be enriched when considering the overall mutation burden. Lastly, 32% of the *PLCB4* mutant cases had more than one mutation in *PLCB4* (*i.e.* potential biallelic mutation), compared to 38% for *NF1*, 14% for *TP53* and 4% for *PTEN* (**Figure 7B**).

**Figure 7.**
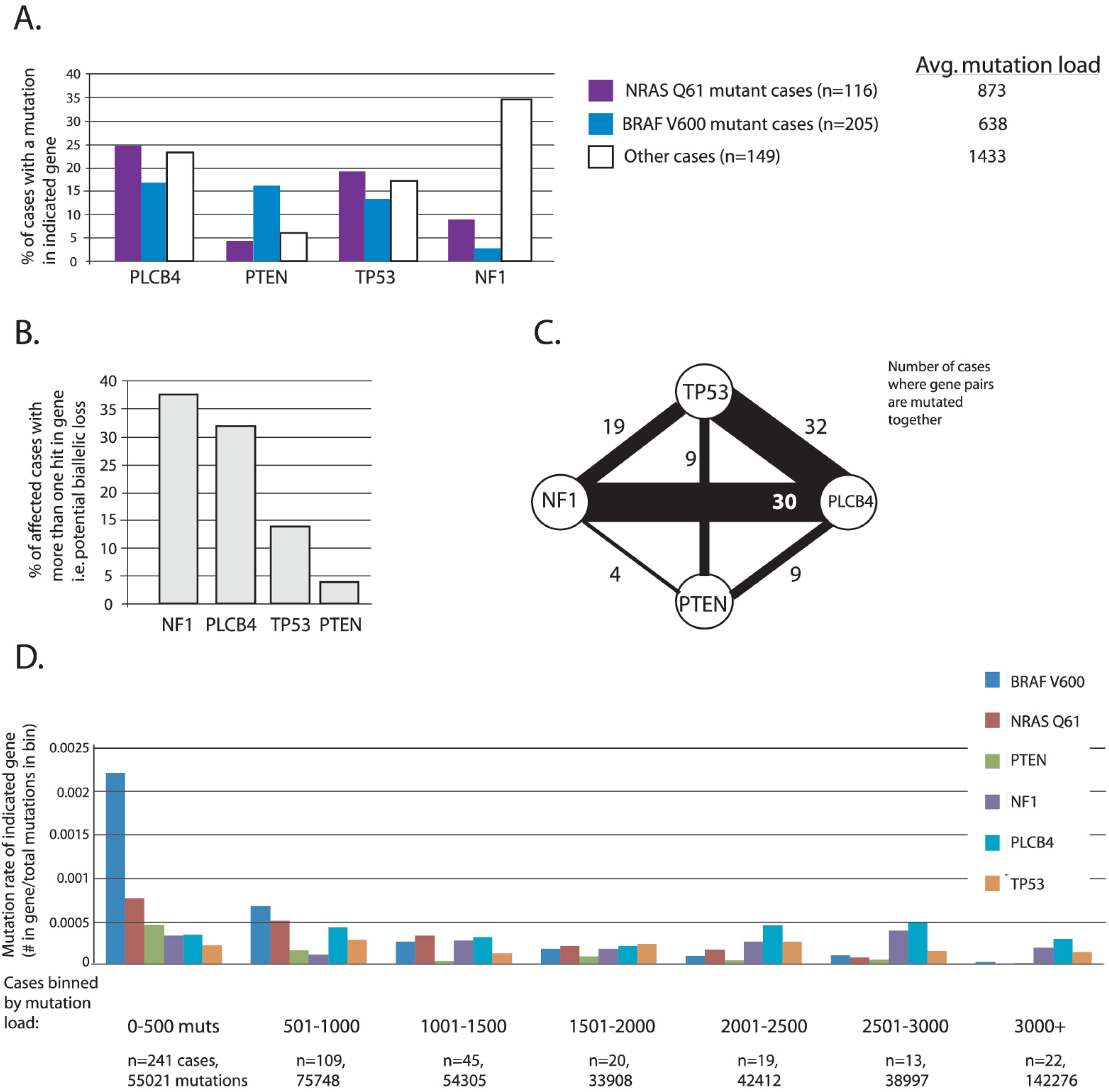
Analysis of PLCB4 mutations in TCGA-SKCM dataset. **A)** Percentage of cases with mutations in PLCB4, PTEN, TP53 or NF1, in cases that were either NRAS-061 or BRAF-V600 mutant, or did not carry these two mutations (’other’). Shown to the right is the average mutation load for NRAS-061, BRAF-V600,or the other cases. **B)** Proportion of cases that had more than one mutation in the indicated gene, out of all cases with a mutation in that gene. **C)** Map illustrating the number of cases with a mutation in more than one of the genes of interest in the study (PLCB4, PTEN, TP53, NF1). PLCB4 mutation was most often found with mutations in TP53 and NF1. **D)** Graph showing the mutation rate of BRAFV600, NRAS061, PTEN, NF1, PLCB4 orTP53 (excluding synonymous mutations) in cases that were binned by their total number of mutations. The number of cases in each bin and the total number of mutations in those cases is shown below the graph.

We then identified all cases that had a mutation in more than one of these four genes of interest (*PTEN*, *NF1*, *PLCB4* and *TP53*) and counted the number of times a double hit for any of the two gene combinations was found (**Figure 7C**). In agreement with the frequencies in Figure 8A, *PTEN* mutations were less often found with other members of this group, particularly *NF1*. *PLCB4* mutations were most often seen with *TP53* and *NF1*, which were found less frequently with each other than with *PLCB4*.

**Figure 8.**
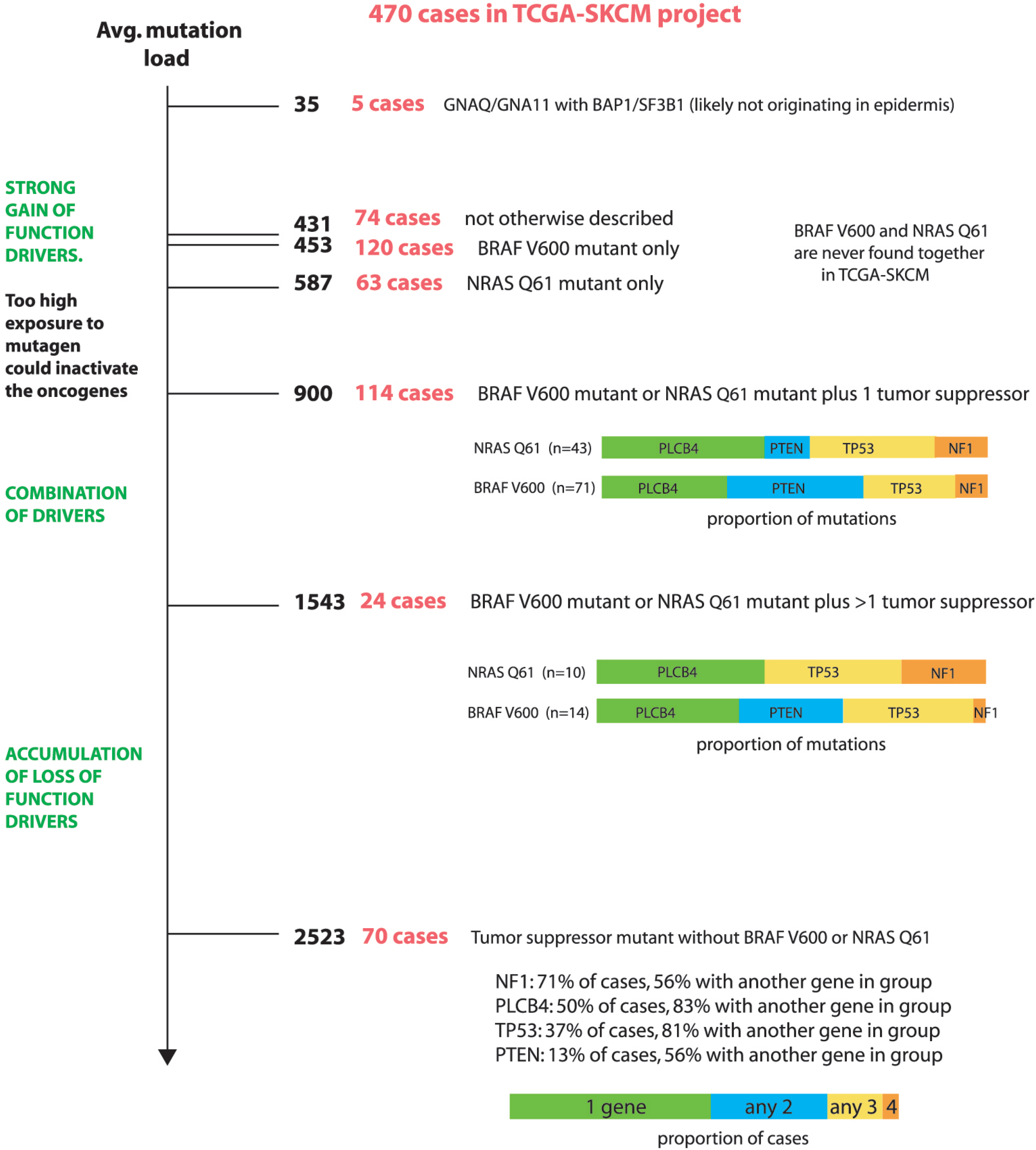
Summary of TCGA-SKCM cases, organized by average mutation load. As mutation load increases, gain of function drivers (BRAF-V600 and NRAS-Q61) become less frequent, while loss of function drivers (NF 1, TP53, PTEN) become more frequent, as do mutations in PLCB4, a candidate tumor suppressor. When mutagen exposure is lower, oncogenic BRAF or NRAS mutations could be more likely to drive melanoma because an inactivating hit to the same allele is less likely. However, when mutagen exposure is higher, it instead allows for the accumulation of loss of function mutations in multiple tumor suppressors. At an intermediate mutation load, BRAF-V600 and NRAS-Q61 mutations co-occur with tumor suppressor mutations. BRAF-V600 preferentially associates with PTEN.

Finally, we grouped all of the cases into bins based upon the total number of small somatic mutations in all genes. This was done to examine the relationship between the overall mutation load with the frequency of mutation in the genes of interest (*BRAF^V6^*^00^, *NRAS^Q^*^61^, *PTEN*, *NF1*, *PLCB4* and *TP53*) (**Figure 7D**). We calculated the mutation rate of *BRAF^V6^*^00^, *NRAS^Q^*^61^, *PTEN*, *NF1*, *PLCB4* and *TP53* in each bin by dividing the total number of small somatic mutations (not including synonymous) found within each gene by the summed total number of mutations in all genes in each bin. The mutation rate of *BRAF^V6^*^00^ and *NRAS^Q61^*was highest in the bin with the smallest mutation load. As mutation load increased, the mutation rate of *BRAF^V6^*^00^ and *NRAS^Q^*^61^ consistently decreased. *PTEN* also generally followed this pattern. In the 2000 to 3000 mutation load/case range, the mutation rate of *BRAF^V6^*^00^, *NRAS^Q^*^61^ and *PTEN* was low, while *NF1* and *PLCB4* reached their maximum. The rate of *NF1* and *PLCB4* mutations declined in the 3000+ mutation load/case bin. *TP53* showed no particular pattern across the bins.

We then organized all of the current TCGA-SKCM cases into **Figure 8**, reflecting on how the overall mutation load relates to mutations in these six genes of interest. At the lowest mutation load (average of 35 total mutations per case), we note that there were five cases of *GNAQ* or *GNA11* oncogenic mutations plus a mutation in either *BAP1* or *SF3B1*. We suspect that these cases could be tumors from uveal or mucosal melanomas, although they were not described as such. Next, there were 74 cases with an average of 431 mutations per case, with no mutations in any of the genes of interest described here. Cases with just *BRAF^V6^*^00^ or *NRAS^Q^*^61^ closely followed, with an average of 453 and 587 mutations per case, respectively. As the mutation load continued to increase, hits in multiple genes of interest accumulated. First, there were cases with just one tumor suppressor mutation plus *BRAF^V6^*^00^ or *NRAS^Q^*^61^ (at 900 mutations/case). The tumor suppressor mutation was more likely to be *PTEN* in the *BRAF^V6^*^00^ mutant subset. Next, there were combinations of two or more tumor suppressor mutations along with either *BRAF^V6^*^00^ or *NRAS^Q^*^61^ (at 1543 mutations/case). Cases with only tumor suppressor mutations made up the last group, which had an average of 2543 mutations per case. Half of these cases had just one mutant tumor suppressor, while the remaining half had more than one mutated (**Figure 8**).

These findings are consistent with previous studies. For example, Mar *et al* found that *BRAF/NRAS* WT cutaneous melanomas have a higher average mutation rate than *BRAF* or *NRAS* mutant melanomas^47^. The mutations in the *BRAF/NRAS* WT tumors were spread over various signaling pathways and included *NF1*, *KIT* and *NOTCH1*^47^. Palmieri *et al* also summarized a number of gene mutations that vary in frequency between *BRAF*/*NRAS* WT vs. mutant melanomas^48^.

The overall pattern is that gain of function drivers are more frequently found when mutation load is lower and loss of function drivers predominate when mutation load is higher. Perhaps when mutagen exposure is low, a stronger acting *BRAF* or *NRAS* mutation can drive the melanoma because an inactivating hit to the same allele is less likely to occur. However, when mutagen exposure is high, it alternatively allows for the accumulation of loss of function mutations in multiple genes, which has a positive synergistic effect when they occur in tumor suppressors.

Our finding that Gα_q/11_ signaling unexpectedly restrains melanocyte growth in the epidermis provides an explanation for the frequent occurrence of *PLCB4* mutations in cutaneous melanoma.

## DISCUSSION

Oncogenic *GNAQ* and *GNA11* mutations are observed at a high frequency in human melanocytic neoplasms in non-epithelial tissues. Still, they are rare in anatomical locations with an epithelial component, such as the inter-follicular epidermis or conjunctiva of the eye. One theoretical explanation for this skewed frequency is that some unknown mechanism mutates *GNAQ* and *GNA11* only in non-epithelial locations. We found that this is not the case. The forced expression of oncogenic GNAQ^Q209L^ in mature melanocytes in mice did not result in cutaneous melanoma, but instead had the opposite effect: the gradual loss of melanocytes from the IFE. Others have hypothesized that subtle geographic variances in the embryonic origin of epithelial versus non-epithelial melanocytes could explain the restriction of certain driver mutations to specific melanoma subtypes^49, 50^. Alternatively, since epidermal melanocytes interact with keratinocytes, whereas non-epidermal melanocytes interact with the mesodermal stromas, it was possible that direct cell contact or paracrine signaling produced by the tissue-specific microenvironment might interfere with the oncogenic signaling pathway^13^. Here, we have presented evidence that the IFE microenvironment drives the loss of melanocytes expressing the GNAQ^Q209L^ oncogene, providing a new model for oncogene specificity.

### Melanocytes taken from the IFE and cultured *in vitro* can switch phenotypes

Previous studies have shown that stable transfection of *GNAQ^Q209L^* induced anchorage- independent growth in soft agar of hTERT/CDK4^R24C^/p53^DD^ mouse melanocytes in a TPA- independent manner, a feature associated with cellular transformation^3, 51^. In addition, GNAQ^Q209L^ signaling produced an abnormal cell morphology with irregular contours. A similar observation was described in *GNAQ^Q209L^*-transformed human melanocytes and melanocytes of a *Gnaq^Q209L^* zebrafish model^52^. Our study complements these previous results to show that melanocytes directly taken from the mouse IFE and grown in primary culture without any other cues survive better if they express *GNAQ^Q209L^*. We also found that *Braf^V600E^* had a stronger effect than *GNAQ^Q209L^* in this same situation, although the promoters driving the expression of these two oncogenes were not the same. In addition, both WT and GNAQ^Q209L^ expressing melanocytes survived better in culture when grown on MEFs. Therefore, the attrition of GNAQ^Q209L^ expressing IFE melanocytes from the mouse tail epidermis is not due to an inherent characteristic in IFE melanocytes because the phenotype is reversible by changing the microenvironment. This is exciting because it suggests that there might be a way to shut the GNAQ^Q209L^ oncogene off through some external cues.

### The IFE microenvironment inhibits melanocytes expressing GNAQ^Q209L^

It is well known that keratinocytes, which make up the vast majority of the epidermis, secrete various growth factors and cytokines in a paracrine manner that regulate growth, survival, adhesion, migration, and differentiation of melanocytes^53^. In WT melanocytes, we found that the presence of dissociated IFE in co-cultures provided a significant boost. In contrast, co-culturing GNAQ^Q209L^ expressing melanocytes with IFE reduced their survival capacity and inhibited cell division. Moreover, the effects of the IFE on melanocyte survival did not require direct cell-cell contact and could be replicated in a transwell culture system. Our studies of IFE melanocytes suggests that GNAQ^Q209L^ expression causes increased cell adhesion, a disorganized actin cytoskeleton and the promotion of long dendrite extensions, which frequently break, possibly related to changes in semaphorin signaling in pathways shared with neurons. We conclude that the microenvironment surrounding melanocytes is the main factor controlling whether GNAQ^Q209L^ signaling is oncogenic or inhibits growth. IFE melanocytes have the innate capacity to be transformed by GNAQ^Q209L^, but whether or not they are permitted to do so is dictated by the microenvironment in which they reside.

### A cellular stress response signature in GNAQ^Q209L^ IFE melanocytes related to vitiligo

Reactive oxygen species (ROS) are highly active radicals produced during multiple cellular processes that, upon accumulation, can damage most biological macromolecules^54^. ROS overproduction in melanocytes by exogenous and endogenous stimuli has been implicated in vitiligo, a common skin depigmentation disorder caused by a loss of melanocytes from the epidermis. ROS accumulation can lead to membrane peroxidation, decreased mitochondrial membrane potential, and apoptosis of melanocytes in vitiligo epidermis^32^. Interestingly, GNAQ^Q209L^ IFE melanocytes shared a similar intracellular response to oxidative stress as observed in vitiligo melanocytes, including activation of p53-dependent signaling and Akt signaling, and induction of both *CDKN2A* (*p16*) and cyclin D1 (*CCND1*) expression^32–34^.

Disruption in these cellular processes might lead to metabolic disturbances, cell cycle arrest, senescence and cell death in melanocytes^32^. Previous studies have shown that melanocytes have an increased susceptibility to oxidative stress compared with keratinocytes or fibroblasts and that *CDKN2A* (*p16*) plays a crucial role in regulating oxidative stress independently of Rb-tumor- suppressor function^55^. Furthermore, oncogene activation in melanocytes generates oxidative stress and increased ROS and p16 expression has been implicated in oncogene-induced melanocyte senescence^56^. Thus, up-regulation of *Cdkn2a* (p16) may reflect a systemic oxidative stress experienced by *GNAQ^Q209L^* melanocytes in an epithelial context.

In support of this idea, Cystatin C (*Cst3*), a potent inhibitor of cysteine proteinases and some lysosomal caspases involved in apoptotic-related processes^37^, was up-regulated in GNAQ^Q209L^ melanocytes. Increased *Cst3* expression was correlated with oxidative stress-induced apoptosis in cultured neurons^57^, and with the induction of intracellular ROS from mitochondria by depletion of glutathione in dendritic cells^58^. Moreover, *S100A8/A9* and *BNIP3* genes involved in cell death and autophagy via ROS-mediated crosstalk between mitochondria and lysosomes were also upregulated^35^. Similarly, we found up-regulation of the pro-apoptotic BOK, an effector of mitochondrial outer membrane permeabilization (MOMP) that triggers membrane permeabilization and apoptosis in the absence of BAX and BAK^36^.

In addition, among the top-ranked DE genes was *Stanniocalcin* (*Stc1*), a glycoprotein hormone involved in calcium/phosphate homeostasis. Significant up-regulation of *STC1* was reported in tumors under hypoxic or oxidative stress^27–31^. Furthermore, *Stc1* expression down-regulates pro-survival Erk1/2 signaling and reduces survival of MEFs under conditions of oxidative stress ^28^.

The slow loss of melanocytes from the IFE in GNAQ^Q209L^ expressing mice is consistent with damage induced by ROS. This represents a novel and intriguing potential consequence of GNAQ^Q209L^ signaling induced by the interfollicular epidermal microenvironment.

### *PLCB4* mutations in cutaneous melanoma

*PLCB4* encodes a protein that binds to Gα_q_ and Gα_11_ and serves as the primary effector of signaling to downstream components in the pathway. If Gα_q/11_ signaling inhibits melanocyte growth in the epidermis, then we wondered whether loss of function mutations in *PLCB4* might promote melanomagenesis. Like Wei *et al.* ^44^, we found that somatic *PLCB4* mutations are frequent in cutaneous melanomas. Excluding synonymous alterations, small somatic mutations in *PLCB4* are present in 97 of the 470 TCGA-SKCM cases (21%). Of the affected cases, most carry a missense mutation, with S670L, G876E and E1104K as the most frequent recurrent substitutions. In addition, 32% of *PLCB4* mutant cases have more than one mutation in the gene (potential biallelic mutation), which was similar to *NF1* at 38% of cases. The pattern of *PLCB4* mutation across the set was not random; instead, *PLCB4* mutations were less frequent in *BRAF^V6^*^00^ mutant cases and more frequent in cases with a higher overall mutation load, alongside *NF1* and *TP53*. Our finding that GNAQ^Q209L^ signaling restrains melanocyte growth in the epidermis provides an explanation for the frequent occurrence of *PLCB4* mutations in cutaneous melanoma. It will be of interest to study the relationship between *PLCB4* loss and frequency and location of metastases, since metastasis involves melanoma cells leaving the epidermal microenvironment and interacting with mesodermal fibroblasts and stromas.

In conclusion, we have shown that the *GNAQ^Q209L^* oncogene switches from promoting to inhibiting melanocyte growth due to paracrine crosstalk with the IFE microenvironment. Our studies revealed evidence for multiple potential mechanisms, including oxidative stress, apoptosis, inhibition of cell division, changes to cell adhesion and cell fragmentation. Hence, the Gα_q/11_ signaling pathway is not only important for uveal melanoma; it also has implications for both vitiligo and cutaneous melanoma. The critical paracrine signal(s) remain to be determined through further biochemical and cell culture studies. It is also possible that a better understanding of the ability of GNAQ^Q209L^ to switch from acting as an oncogene to an inhibitor of melanocyte growth could lead to new therapies for uveal melanoma, which lacks effective treatment options for metastatic disease.

## METHODS

### Mice

Animal research was conducted under the approval of the UBC Animal Care Committee (Protocols A18-0080 and A19-0148, C.D.V.R.). DNA from ear notches was isolated using DNeasy columns (Qiagen) and amplified using PCR. *Mitf-cre* (Tg(Mitf-cre)7114Gsb), *Tyrosinase-creERT^2^* (Tg(Tyr-cre/ERT2)13Bos/J), *Rosa26-LoxP-Stop-LoxP-GNAQ^Q209L^* (Gt(ROSA)26Sor^tm1(GNAQ*)Cvr^), *Rosa26-LoxP-Stop-LoxP-tdTomato* (Gt(ROSA)26Sor^tm14(CAG-tdTomato)Hze^), *Rosa26-LoxP-Stop-LoxP-LacZ* (Gt(Rosa)26Sor^tm1Sor^/J), and *Braf^CA^* (*Braf^tm1Mmcm^*) mice were genotyped as previously described^14–17, 20, 59^. All strains were backcrossed to the C3HeB/FeJ genetic background for at least 3 generations.

### LacZ staining

We crossed *R26-fs-GNAQ^Q209L^*/+; +/+ mice to *R26-fs-LacZ/+*; *Tyr-creERT^2^/+* mice and genotyped the resulting progeny. At 4 weeks of age, two groups of mice were tamoxifen-treated for 5 consecutive days: *R26-fs-GNAQ^Q209L^*/*R26-fs-LacZ*; *Tyr-creER/+* (GNAQ-LacZ) mice and +/*R26-fs-LacZ* mice; *Tyr-creER/+* (WT-LacZ) mice. Tamoxifen treatment consisted of one daily intraperitoneal injection of 1 mg tamoxifen (Sigma T5648) alongside a topical treatment for the tail skin (tails were dipped for 5 seconds in 25 mg/ml 4-hydroxytamoxifen (Sigma H6278) in DMSO). Mice were harvested either 1 week or 8 weeks after tamoxifen treatment. At each experimental endpoint, a piece of tail skin of 1.5 cm in length was incubated in X-gal solution for 48 hours as previously described^60, 61^. The tail dermis and epidermis were split using sodium bromide and fixed in 4% PFA. The number of LacZ-positive cells was counted in three rows of epidermal scales per sample.

### Histochemistry

For H&E staining, mouse tail skin samples were fixed in 10% buffered formalin overnight at room temperature with gentle shaking, dehydrated, cleared, and embedded in paraffin. 5 μm sections were taken for H&E staining using standard methods and imaged using a DMI 6000B microscope (Leica). To examine tdTomato expression in sections, samples were fixed in 10% buffered formalin overnight at 4°C, taken through a sucrose gradient, embedded in O.C.T., and sectioned at 10 μm. After washing in PBS for 30 minutes, sections were counter stained with DAPI and imaged using a Histotech III slide scanner.

### Isolation of inter-follicular melanocytes from mouse tail skin

We crossed *R26-fs-GNAQ^Q209L^/+* mice to *+/R26-fs-tdTomato; Mitf-cre/+* mice and identified tdTomato expressing progeny by genotyping. At four weeks of age, the tails were waxed to remove hair follicles. The next day, tail skin from GNAQ^Q209L^ and WT tdTomato-positive mice was harvested, and the IFE was split from the dermis by gentle dispase treatment. The IFE was then incubated with trypsin into single cells, using forceps to scrape cells from the scales to accelerate the process. (A detailed description of this method will be concurrently submitted to *Bio Protocol*).Cells were centrifuged and the pellet was resuspended in suspension buffer (HBSS + 10% FBS + EDTA 0.1mM EDTA) prior to FACS sorting.

### FACS

Dissociated cells were first analyzed for viability based on forward and side scatter plots and using eBioscience™ Fixable Viability Dye. Next, filters for forward scatter (height versus width) and side scatter (height versus width) were used to select single cells. Lastly, viability dye fluorescence (y-axis) was plotted against tdTomato (x-axis) and gates were set to capture only viable tdTomato positive cells. Cells from mice lacking *Mitf-cre* (*+/R26-fs-tdTomato; +/+*) were used as negative controls to define the threshold for tdTomato-positive cells. Sorted cells were collected in 15ul of HBSS+FBS 10% +EDTA 0.1 mM solution and 1 ul of RiboLock RNase inhibitor (Thermo Scientific, EO0381). For Annexin V staining, we used the Annexin V Fluos staining kit according to the manufacturer’s directions (Sigma Aldrich, 11858777001).

### Primary cell culture

Cells were plated into 96-well plates previously coated with 0.1 mg/ml fibronectin in Dulbecco’s Modified Eagle’s Medium (DMEM) supplemented with 10% Fetal Bovine Serum, 1% penicillin-streptomycin (15140122 Thermo-Fisher), 2 mM L-Glutamine, and 1 mM Sodium Pyruvate. For cultures of sorted melanocytes only, cells were plated at 6000 to 8000 cells per well. For co-culture with MEFs, 6000 to 8000 FACS sorted melanocytes were plated into 96-well plates previously seeded with MEFs that had formed a confluent monolayer. For direct co-culture experiments with IFE, the IFE was dissociated and plated at 100,000 cells per well (*i.e.* with no FACS sorting for the melanocytes first). For transwell co-culture with IFE, 6000 to 8000 FACS sorted melanocytes were plated in the well suspended above 100,000 dissociated IFE cells plated below. All cultures were incubated at 37°C in 5% CO_2_. Media was changed every third day by exchanging 1/3 of the existing volume. Images were taken at 5x, 10x, and 20x magnification. tdTomato positive cells in the 5x field of view were counted as live melanocytes.

### Live cell imaging

Sorted tdTomato positive melanocytes or IFE cells containing tdTomato positive melanocytes were plated on 0.1 mg/ml fibronectin-coated coverslips within a 35mm cell culture dish. Cells were incubated overnight before imaging in a live-cell imaging culture chamber. Cells were imaged at 37°C (5% CO_2_) every 5 minutes for 20 hours at 10X magnification. Cell movements were determined using the MTrackJ software on ImageJ and analysis was performed using Chemotaxis and Migration Tool (Ibidi).

### Automated image analysis

Custom Matlab scripts were utilized to analyze cell morphology and actin fiber orientation in fixed cells as described in Haage *et al*^62^ and posted to GitHub. Cell contours were obtained automatically by processing confocal z-projections of cells stained for F-actin (Phalloidin) in MatLab. Images were first blurred with a Gaussian filter, and then an edge detection algorithm was applied to identify cell borders. The resultant binary images were refined through successive dilations and erosions to yield the final cell contour. These contours were used to measure cell area, and circularity (4π*area/perimeter^2^), protrusions. The number and length of protrusions were quantified automatically in MatLab. First, cell contours were identified as outlined above. Next, we obtained the contour coordinates of the convex hull of the binary image representing the cell area. At each point along the cell contour, we computed the minimum distance between the convex hull and the actual cell contour. Based on these distances, minima corresponding to protrusions could be identified. To be counted as protrusions, minima had to be at least 10 pixels apart along the contour and of height greater than 5 pixels. Based on the coordinates of adjacent peaks, the width, height, and aspect ratio of protrusions could be computed.

### Actin fiber organization

First, we identified cell contours as described above. Next, the cell was subdivided into 32x32 pixel windows overlapping by 50%. We then computed the two-dimensional Fourier transform of each window. If a window contains no fibers, the Fourier transform will be a central, diffuse point of bright pixels. However, if a window contains aligned fibers, the Fourier transform will consist of an elongated accumulation of bright pixels at a 90 degree angle to the original fibers. Based on the aspect ratio and orientation of the Fourier transform, we determined fibrousness and fiber orientation in a given window. The data for individual windows could then be compared across the entire cell to estimate the cell fibrousness, defined here as the percentage of cell area (% of windows) with aspect ratio greater than a cut-off value.

### RNA-seq data and bioinformatics analysis

Total RNA from sorted cells was extracted using Trizol (Life Technologies) following the manufacturer’s protocol. For each library preparation, sorted melanocytes from two mice of the same litter were pooled to increase the number of cells for RNA extraction. We generated 3 Wt libraries and 3 GNAQ libraries from a total of 12 mice. Sample quality control was performed using the Agilent 2100 Bioanalyzer. Qualifying samples (RNA Integrity Number ≥ 9) were then prepped following the standard protocol for the NEB next Ultra ii Stranded mRNA (New England Biolabs). Sequencing was performed on the Illumina NextSeq 500 with Paired End 42bp × 42bp reads. Sequencing data was demultiplexed using Illumina’s bcl2fastq2. De-multiplexed read sequences were then aligned to the *Mus Musculus* mm10 reference sequence using STAR aligner^63^. The fastq sequences have been deposited at the Sequencing Read Archive (SRA) of the NCBI under BioProjectID PRJNA736153.

[Private link to PRJNA736153 sequences: https://dataview.ncbi.nlm.nih.gov/object/PRJNA736153?reviewer=no8fe0hmv2ukgvoplvlu0kbu bu]

Assembly and differential expression analysis were performed using Cufflinks^64^ through bioinformatics apps available on Illumina Sequence Hub. Gene Ontology and KEGG Pathways analysis was performed using DAVID Bioinformatics Resources 6.8^65, 66^. Gene Set Enrichment Analysis (GSEA) was performed using the JAVA GSEA 2.0 program^67^. The gene sets used for analysis were the Broad Molecular Signatures Database gene sets H (Hallmark gene sets).

## Statistical analysis

Sample-size estimation: Target sample sizes were not explicitly computed during the study design phase because the standard deviations between wildtype and GNAQ^Q209L^ epidermal melanocytes were unknown. However, in our past experience of studying skin pigmentation phenotypes on inbred genetic backgrounds, 4 mice of each genotype is usually sufficient to detect statistically significant differences. For primary cell culture experiments, we planned to perform each experiment in 3 complete and independent runs, from FACS collection to cell culture. For RNAseq, we planned to generate 3 wildtype libraries and 3 GNAQ^Q209L^ libraries, based on previously published studies.

Replicates: All replicates were biological replicates, *i.e.* each sample or independently derived cell culture was established from a different mouse. In some experiments, one biological replicate contained melanocytes collected from two mice of the same age and genotype, which were pooled together to form one sample. When pooling occurred, it was done at the FACS step. The number of replicates (number of mice and number of melanocytes derived from the mice) can be found in Supplementary Table 10, organized by figure number.

Statistical reporting: The statistical tests that were used in each figure are described and shown in Supplementary Tables 10 and 11. Raw data was plotted in graphs. Except for the survival graphs, the mean for each group and the standard error of the mean is shown. Exact p values, t values, degrees of freedom and number of replicates are detailed in Supplementary Tables 10 and 11. Statistical analysis was performed using GraphPad Prism software. For RNAseq, the magnitude and significance of differential gene expression and multiple test corrections were determined using the Cufflinks suite via the Illumina Sequence Hub. Gene Ontology and KEGG Pathways analysis was performed using DAVID Bioinformatics Resources 6. Gene Set Enrichment Analysis (GSEA) was performed using the JAVA GSEA 2.0 program.

Group allocation: Samples were allocated into experimental groups based on genotype (WT or GNAQ) or culture condition. WT and GNAQ mice/melanocytes that were compared to each other were produced from the same set of breeder parents. Masking was used whenever possible (melanocytes/skin samples with an obvious difference in pigmentation are not maskable).

## ACKNOWLEDGEMENTS

We thank Dr. Gregory Barsh for contributing the *Mitf-cre* mouse line. Research was supported by grants from the Canadian Institutes for Health Research (grant MOP-79511 to C.D.V.R and grant PJT-168868 to G.T.).

## AUTHOR CONTRIBUTIONS

O.U. and C.D.V.R conceived the project and designed the experiments. O.U. and A.H. performed experiments. O.U., A.H., G.T. and C.D.V.R. performed data analyses. O.U. and C.D.V.R wrote the manuscript.

## Index of Supplementary Files

**Supplementary Table 1:** Genes with FPKM > 0.1 in WT and/or GNAQ Q209L melanocytes.

**Supplementary Table 2.** Top 20 most highly expressed genes in mouse WT IFE melanocytes.

| Rank | Gene symbol | Gene name | Directly related to pigment production/melanosome biology? | Reference |
| --- | --- | --- | --- | --- |
| 1 | Pmel | Premelanosome protein | Yes | (Lee et al., 1996) |
| 2 | Dct | Dopachrome tautomerase | Yes | (Tsukamoto et al., 1992) |
| 3 | Mir5133 | MicroRNA 5133 |  |  |
| 4 | Tyrp1 | Tyrosinase-related protein 1 | Yes | (Kobayashi et al., 1994) |
| 5 | Ptgds | Prostaglandin D2 synthase (brain) |  |  |
| 6 | Mlana | Melan-A | Yes | (Du et al., 2003) |
| 7 | Eef1a1 | Eukaryotic translation elongation factor 1 alpha 1 |  |  |
| 8 | Vim | Vimentin |  |  |
| 9 | Cd63 | CD63 antigen | Yes | (van Niel et al., 2011) |
| 10 | Krt14 | Keratin 14 |  |  |
| 11 | Lgals1 | Lectin, galactose binding, soluble 1 |  |  |
| 12 | Fth1 | Ferritin heavy polypeptide 1 |  |  |
| 13 | Rplp1 | Ribosomal protein, large, P1 |  |  |
| 14 | Gstp1 | Glutathione S-transferase, pi 1 |  |  |
| 15 | Slc24a5 | Solute carrier family 24, member 5 | Yes | (Lamason et al., 2005) |
| 16 | Gpnmb | Glycoprotein (transmembrane) nmb | Yes | (Loftus et al., 2009) |
| 17 | Rps27rt | Ribosomal protein, S27, retrogene |  |  |
| 18 | Ftl1 | Ferritin light polypeptide 1 |  |  |
| 19 | Krt10 | Keratin 10 |  |  |
| 20 | Rpl8 | Ribosomal protein L8 |  |  |

**Supplementary Table 3.** Differentially expressed genes down-regulated in GNAQ-Q209L IFE melanocytes, sorted by z_score.

**Supplementary Table 4.** Differentially expressed genes up-regulated in GNAQ-Q209L IFE melanocytes, sorted by Z_score.

**Supplementary Table 5.**
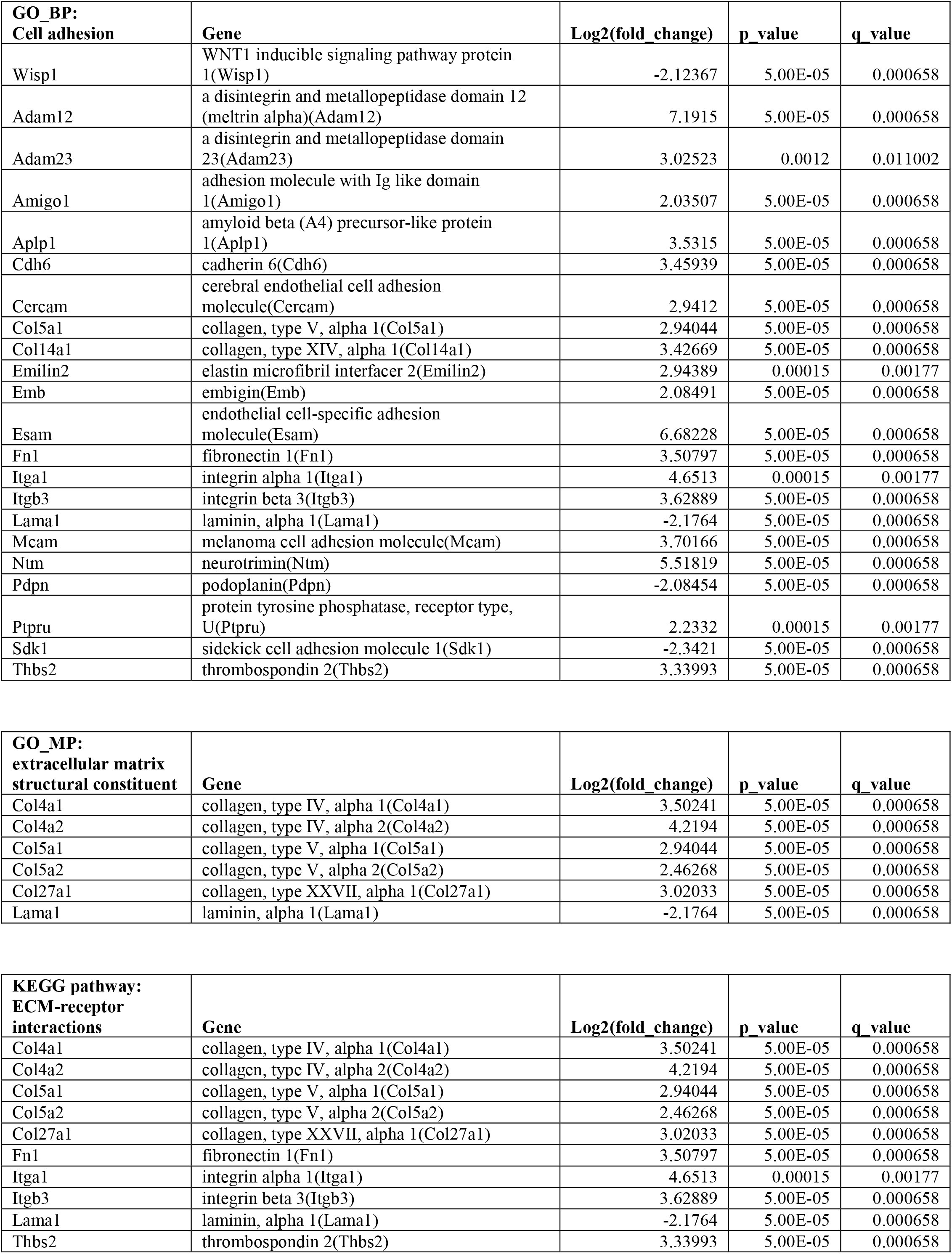
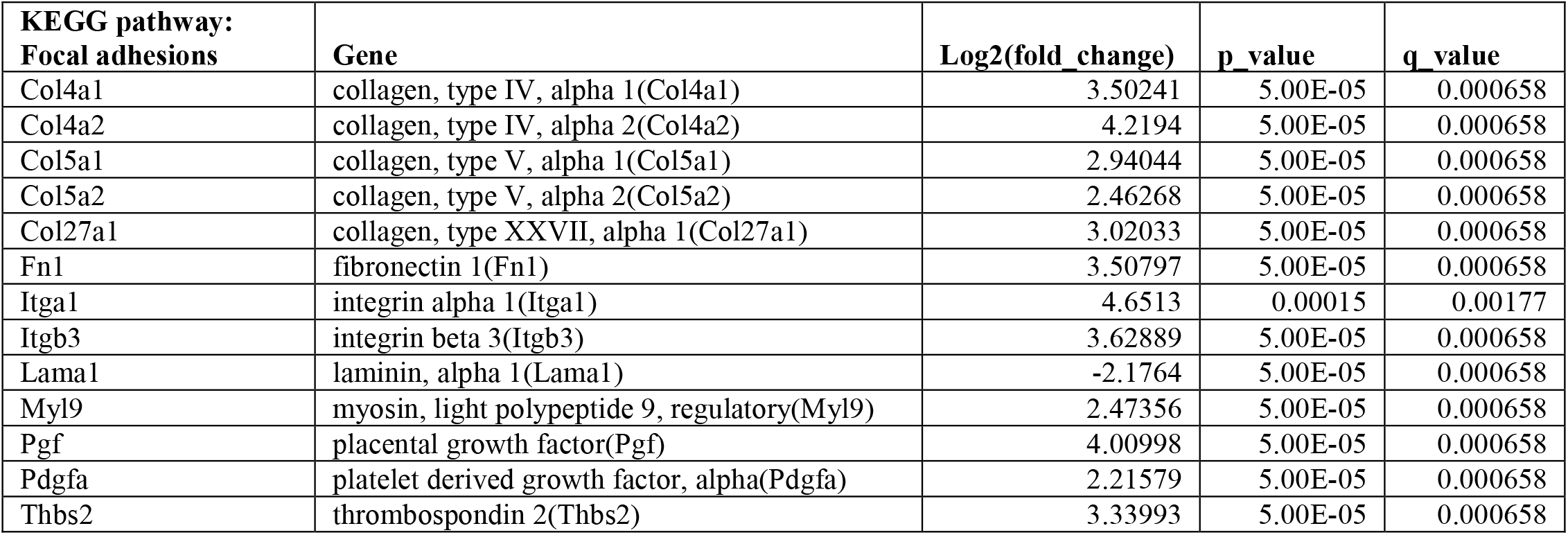
Genes supporting pathway analysis terms related to cell adhesion, focal adhesion and the extracellular matrix.

**Supplementary Table 6.**
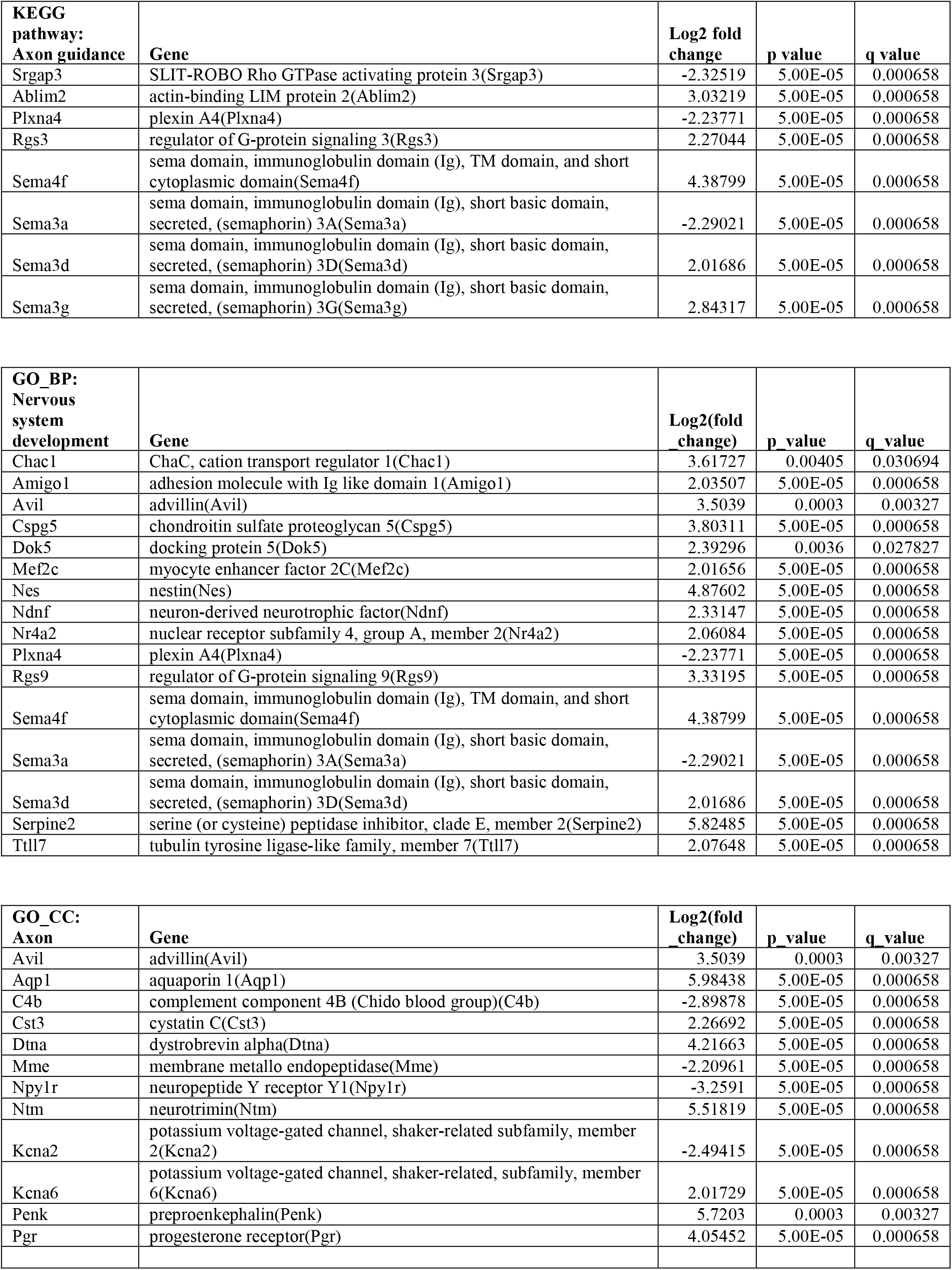
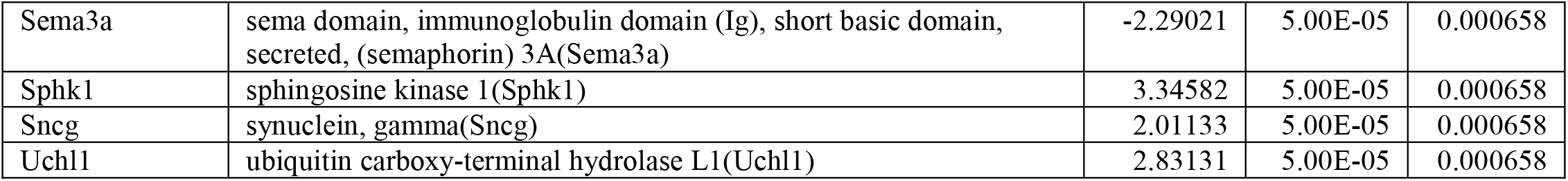
Genes supporting pathway analysis terms related to axon guidance, nervous system development, and axon cellular component.

**Supplementary Table 7.** Identification of *GNAQ* hotspot mutations among human malignant melanomas potentially arising in the epidermis. The Cosmic database was searched for GNAQ Q209 or R183 missense mutations in tumors with the following criteria: <u>Primary site</u>: Skin and <u>Histology</u>: Malignant melanoma. Must have one of the following terms for <u>Subhistology</u>: Superficial spreading, Nodular, Spitzoid, Lentigo maligna, Acral lentiginous, Amelanotic, Epithelioid, Nevoid, or NS (Not Specified). (Excluded subhistology terms: Benign, Desmoplastic, Mucosal, Blue). Must have one of the following terms for <u>Subsite</u>: ear, lip, elbow, back, upper back, lower back, ankle, trunk, groin, hand, knee, chest, scalp, face, leg, shoulder, arm, breast, neck, flank, extremity, upper arm, forearm, foot, chronically sun exposed, intermittently sun exposed, eye, non chronically sun exposed or NS (Not specified). (Excluded subsite terms: mucosal, axilla, subungual, penis, nipple, vulva). The total number of *GNAQ* tested samples defined by these terms was 2753. Samples that could have arisen in the epidermis are highlighted in yellow. N/A, non-applicable; NGS, Next generation sequencing.

| Cosmic Sample ID | PMID | GNAQ mutation | Other key mutations | Scope of study |
| --- | --- | --- | --- | --- |
| 1175905 | 19078957 | Q209L | N/A | GNAQ only |
| 2543893 | 26275246:<br>"Malignant melanoma arising in blue nevus" | Q209L | N/A | N/A |
| 2724266 | 28481359 | Q209H (frq =0.04) | N/A | N/A |
| 2719694 | 28481359 | Q209L | BAP1 | MSK-Impact screen |
| 2718695 | 28481359 | Q209L | SF3B1<sup>R625C</sup> | MSK-Impact screen |
| 1993392 | 21726664 | Q209L | None | 30 gene screen |
| 1993559 | 21726664 | Q209P | None | 30 gene screen |
| 2088283-2088289<br>From one patient | 24504448 | Q209P | BRAF<sup>V600E</sup> | NGS |
| 2380390 (RP-A690) | 26091043 | Q209P | SF3B1<sup>R625H</sup> | NGS (TCGA-SKCM) |
| 2121734 (ER-A2NF) | 26091043 | Q209P | SF3B1<sup>R625H</sup> | NGS (TCGA-SKCM) |
| 2380420 (BF-AAP8) | 26091043 | Q209P | BAP1 | NGS (TCGA-SKCM) |

**Supplementary Table 8.** Identification of *GNA11* hotspot mutations among human malignant melanomas potentially arising in the epidermis. The Cosmic database was searched for GNA11 Q209 or R183 missense mutations in tumors with the following criteria: <u>Primary site</u>: Skin and <u>Histology</u>: Malignant melanoma. Must have one of the following terms for <u>Subhistology</u>: Superficial spreading, Nodular, Spitzoid, Lentigo maligna, Acral lentiginous, Amelanotic, Epithelioid, Nevoid, or NS (Not Specified). (Excluded subhistology terms: Benign, Desmoplastic, Mucosal, Blue). Must have one of the following terms for <u>Subsite</u>: ear, lip, elbow, back, upper back, lower back, ankle, trunk, groin, hand, knee, chest, scalp, face, leg, shoulder, arm, breast, neck, flank, extremity, upper arm, forearm, foot, chronically sun exposed, intermittently sun exposed, eye, non chronically sun exposed or NS (Not specified). (Excluded subsite terms: mucosal, axilla, subungual, penis, nipple, vulva). The total number of *GNA11* tested samples defined by these terms was 2295. Samples that could have arisen in the epidermis are highlighted in yellow. N/A, non-applicable; NGS, Next generation sequencing.

| Cosmic Sample ID | PMID | GNA11 mutation | Other mutations | Scope of study |
| --- | --- | --- | --- | --- |
| 2544691 | 26825879 | Q209L | None | 4 genes screened |
| 1838375 | 22817889:<br>"uveal melanoma" | Q209L | N/A | N/A |
| 2013581 | 22842228:<br>"uveal melanoma" | R183C | N/A | N/A |
| 2013602 | 22842228 | R183C | NRAS<sup>Q61H</sup> | NGS exome |
| 2013703 | 22842228:<br>"uveal melanoma" | Q209L | N/A | N/A |
| 2013594 | 22842228 | Q209L | EIF1AX | NGS exome |
| 753596 | none | R183C | N/A - Cell line | N/A |
| 2237811 | 24714776:<br>"most likely represents metastatic uveal melanoma" | Q209L | N/A | N/A |
| 2380396 (RP-A6K9) | 26091043 | Q209L | BAP1 | NGS (TCGA-SKCM) |
| 2339744 (W3-A285) | 26091043 | R183C | NRAS<sup>Q61R</sup> | NGS (TCGA-SKCM) |
| 2121738 (ER-A3ET) | 26091043 | Q209L | P53<sup>Y220C</sup>, PLCB4 splice | NGS (TCGA-SKCM) |
| 2121737 (ER-A3ES) | 26091043 | Q209L | SF3B1<sup>R625H</sup> | NGS (TCGA-SKCM) |

**Supplementary Table 9.** SSM mutation data from 470 TCGA-SKCM cases, along with total number of mutations and primary diagnosis.

**Supplementary Table 10.** Details on statistical tests.

|  | **Comparison of:** | **Statistical test used:** | **Exact p value:** | **Other values:** | **Degrees of freedom:** | **Definition of replicates:** |
| --- | --- | --- | --- | --- | --- | --- |
| **1B** | Average number of LacZ+ cells per scale in Wt vs GNAQ mice at 1 week of age | Unpaired t-test (two tailed) | 0.18 | t=1.491 | df=7 | Experiment performed on 4 Wt mice and 5 GNAQ mice.<br>At least 25 scales were examined per mouse (depending on the number of scales in the row). |
| **1B** | Average number of LacZ+ cells per scale in Wt vs GNAQ mice at 8 weeks of age | Unpaired t-test (two tailed) | 0.045 | t=2.367 | df=8 | Experiment performed on 5 Wt mice and 5 GNAQ mice.<br>At least 25 scales were examined per mouse (depending on the number of scales in the row). |
| **1D** | Percentage of scales affected by loss of boundaries at 8 weeks of age | Kolmogorov-Smirnov test | 0.18 |  |  | Experiment performed on 5 Wt mice and 5 GNAQ mice.<br>At least 25 scales were examined per mouse (depending on the number of scales in the row). |
| **1D** | Percentage of scales affected by loss of melanin at 8 weeks of age | Kolmogorov-Smirnov test | 0.048 |  |  | Experiment performed on 5 Wt mice and 5 GNAQ mice.<br>At least 25 scales were examined per mouse (depending on the number of scales in the row). |
| **2B** | Percentage of tomato positive cells sorted from Wt vs GNAQ IFE | Unpaired t-test (two tailed) | 0.012 | t=3.132 | df=9 | Percentages measured in 6 Wt tails/FACS runs and 5 GNAQ tails/FACS runs |
| **2C** | Survival of WT, GNAQ and BRAF IFE melanocytes plated on fibronectin | 2 way ANOVA with Tukey's multiple comparisons test | $3.3 \times 10^{-5}$ for genotype<br>$1.6 \times 10^{-5}$ for time | Geisser-Greenhouse's epsilon = 0.3539 | | Fraction of survival measured in 10 independent cell cultures derived from 3 or 4 mice of each genotype |
| **2H** | Survival of WT vs GNAQ IFE melanocytes plated on MEFs | 2 way ANOVA with Tukey's multiple comparisons test | 0.48 for genotype<br>0.31 for time | Geisser-Greenhouse's epsilon = 0.3030 |  | Fraction of survival measured in 3 and 2 independent cell cultures derived from 3 GNAQ mice and 2 WT mice |
| **3A** | Survival of WT melanocytes plated with IFE vs WT melanocytes plated on fibronectin | 2 way ANOVA | 0.0071<br>for culture condition<br>$1.0 \times 10^{-5}$<br>for time | Geisser-Greenhouse's epsilon = 0.3081 | | Fraction of survival measured in 3 and 3 independent cell cultures derived from 6 Wt mice. |
| **3B** | Survival of GNAQ melanocytes plated with IFE vs GNAQ melanocytes plated on fibronectin | 2 way ANOVA | 0.0053<br>for culture condition<br>0.00063<br>for time | Geisser-Greenhouse's epsilon = 0.3370 |  | Fraction of survival measured in 3 and 3 independent cell cultures derived from 6 GNAQ mice. |
| **3D** | Circularity of WT vs GNAQ melanocytes plated with IFE | Unpaired t-test (two tailed) | 0.000000050 | t=5.675 | df=193 | Averages calculated from 116 WT cells and 79 GNAQ cells from four independent culture experiments |
| **3E** | Protrusion length of WT vs GNAQ melanocytes plated with IFE | Unpaired t-test (two tailed) | 0.00041 | t=3.593 | df=207 | Averages calculated from 115 WT cells and 94 GNAQ cells from four independent culture experiments |
| **3F** | Number of protrusions in WT vs GNAQ melanocytes plated with IFE | Unpaired t-test (two tailed) | 0.0096 | t=2.615 | df=207 | Averages calculated from 115 WT cells and 94 GNAQ cells from four independent culture experiments |
| **3G** | Cell area of WT vs GNAQ melanocytes plated with IFE | Unpaired t-test (two tailed) | $2.6 \times 10^{-13}$ | t=7.862 | df=193 | Averages calculated from 116 WT cells and 79 GNAQ cells from four independent culture experiments |
| **3I** | Percentage of melanocytes cultured with IFE exhibiting fragmentation of dendrites | Unpaired t-test (two tailed) | 0.021 | t=2.875 | df=8 | Percentage calculated from 5 WT cultures and 5 GNAQ cultures (total cells tracked = 187 and 98) |
| **4B** | Total distance traveled by WT vs GNAQ cells co-cultured with IFE | Unpaired t-test (two tailed) | 0.86 | t=0.1827 | df=114 | Averages calculated from 57 WT cells and 59 GNAQ cells from four independent culture experiments |
| **4C** | Directness of cell migration in WT vs GNAQ cells co-cultured with IFE | Unpaired t-test (two tailed) | 0.18 | t=1.359 | df=106 | Averages calculated from 59 WT cells and 49 GNAQ cells from four independent culture experiments |
| **4D** | Percentage of cells that divided during 20 hours observation, when co-cultured with IFE. | Ordinary 1 way ANOVA with Tukey's multiple comparisons test | 0.0040 overall |  |  | Averages calculated from 5 WT, 5 GNAQ and 3 BRAF independent cultures. Total cell number tracked: 187 WT cells, 98 GNAQ cells and 121 BRAF cells. |
| **4H** | Average time for cytokinesis in cells plated with IFE. | Ordinary 1 way ANOVA, with Tukey's multiple comparisons test | $3.0 \times 10^{-10}$ overall | | | Average time calculated from 1 GNAQ, 11 WT and 14 BRAF cells that divided in 5, 5 or 3 independent culture experiments. |
| **5A** | Survival of WT melanocytes directly co-cultured with IFE vs transwell cultured with IFE | 2 way ANOVA | 0.022 for culture condition |  |  | Fraction of survival calculated from 3 and 3 independent cell cultures for each condition, derived from 6 WT mice. |
| **5B** | Survival of GNAQ melanocytes directly co-cultured with IFE vs transwell cultured with IFE | 2 way ANOVA | 0.27 for culture condition |  |  | Fraction of survival calculated from 3 and 3 independent cell cultures for each condition, derived from 6 GNAQ mice. |
| **6C** | Cell fibrousness in WT vs GNAQ melanocytes plated with IFE | Unpaired t-test (two tailed) | $1.8 \times 10^{-12}$ | t=7.544 | df=193 | Averages calculated from 116 WT cells and 79 GNAQ cells from four independent culture experiments |

**Supplementary Table 11.** Data used to create graphs and calculate statistics in Figures 1-6.

**Video 1.** WT melanocytes co-cultured with IFE, 625 minute time lapse.

**Video 2.** GNAQ melanocytes co-cultured with IFE, 625 minute time lapse.

